# PandaMap: A Python Package for Comprehensive Visualization of Protein–Ligand Interaction Networks

**DOI:** 10.64898/2026.08.06.743421

**Authors:** Pritam Kumar Panda

## Abstract

Protein–ligand interaction diagrams are a routine part of structural and medicinal chemistry, but the tools that produce them tend to force a choice: comprehensive detection with tabular output, publication-quality figures behind a licence, or a scripting environment that assumes expertise. **PandaMap** (**P**rotein **AND** lig**A**nd interaction **MAP**per) is an open-source Python package that produces a 2D interaction diagram, an interactive 3D viewer, a text report, a machine-readable CSV, and a four-panel graphical summary from a single command. It reads PDB, mmCIF and PDBQT files, detects 15 interaction classes using crystallographically validated distance thresholds, and depends only on NumPy, Matplotlib, BioPython and Requests; RDKit improves the 2D ligand layout when present but is not required. Hydrogen bonds are filtered on the true D–H*· · ·* A angle when the structure contains explicit hydrogens, matching PLIP’s 100◦ criterion on the same evidence, and on distance alone otherwise, with the provenance of each measurement recorded. We benchmarked the package on three complexes chosen for different chemistry: enolase with a phosphonate transition-state analogue (PDB 1ELS), the EGFR kinase with erlotinib (1M17), and aldose reductase with IDD594 (1US0). PandaMap recovers the contacts these structures are known for, including the EGFR hinge hydrogen bond to MET769 and the IDD594 bromine*· · ·* THR113 halogen bond, both at distances identical to PLIP’s. All detection thresholds, scoring weights and the exact commands used are given in the Supplementary Information, and the release carries a regression suite covering each interaction class. PandaMap 4.3.0 is available on PyPI under the MIT licence.

## 1 Introduction

Anyone who has prepared a figure showing how a ligand sits in its binding site knows the routine: run a detection tool, get a table of contacts, then redraw the whole thing by hand because the output is not something you would put in a paper. The Protein Data Bank now holds over 230,000 structures [1], and the number of these diagrams being drawn has grown with it, but the tooling has stayed oddly divided.

PLIP [2, 3] detects interactions carefully and is the closest thing the field has to a reference implementation, but its primary output is tabular and it depends on external OpenBabel and DSSP installations. LigPlot+ [4] produces the schematic diagrams that appear in a great many papers, though it is licensed and offers limited control over the result. PyMOL and ChimeraX will render anything you can describe, provided you are willing to write the script. MDAnalysis and ProLIF compute interaction fingerprints across trajectories, which is a different problem from drawing one good figure of one frame.

PandaMap is an attempt to cover the ordinary case well: one command, one structure file, and a figure you can use, plus the underlying numbers in a form you can process. It parses PDB, mmCIF and PDBQT, picks the ligand automatically or on request, detects 15 classes of contact using published distance thresholds, and writes a 2D diagram, an interactive 3D viewer, a text report, a CSV of every contact, and a four-panel graphical summary. The mandatory dependencies are NumPy, Matplotlib and BioPython. RDKit gives a better 2D ligand layout when installed, and the package works without it.

This paper documents the package and benchmarks it against PLIP on three complexes chosen for different chemistry. Users upgrading from 4.2.x should note that the behaviour of several interaction classes changed in 4.3.0; the differences are set out in Section 5.

## 2 Design and Implementation

### 2.1 Software Architecture

PandaMap is written entirely in Python 3 (*≥*3.7) and organised into four principal modules within the `pandamap` namespace (Figure 1). The core module (core.py) contains three main classes: `MultiFormatParser`, which wraps BioPython [5] `Structure` objects and handles PDBQT-to-PDB conversion; `SimpleLigandStructure`, which owns the 2D projection, ring detection, and bond topology pipeline; and `HybridProtLigMapper`, the primary analysis orchestrator. A second module, `improved_interaction_detection.py`, post-processes raw interaction lists through residue-level deduplication and PLIP-like threshold enforcement before generating formatted text reports. The `create_3d_view.py` module assembles a self-contained HTML page embedding the structure as a base64-encoded PDB string within a 3Dmol.js [6] viewer template. Finally, `cli.py` exposes a `pandamap` entry point via `argparse` that drives the end-to-end pipeline from the terminal.

**Figure 1:**
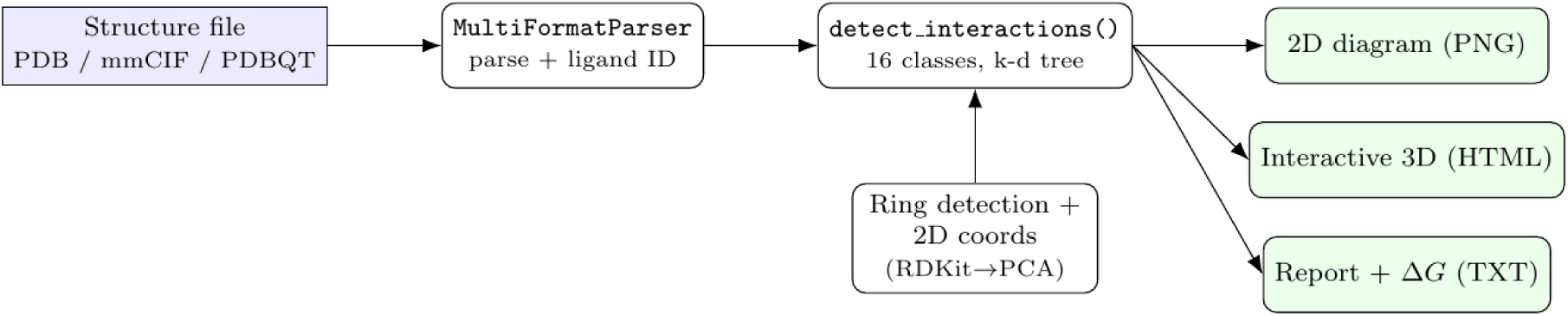
PandaMap architecture and data flow. A single structure file is parsed, the ligand is identified, interactions across 15 classes are detected with a BioPython `NeighborSearch` k-d tree, and three complementary outputs are produced.

Mandatory runtime dependencies are deliberately minimal: NumPy (*≥*1.20) [7] for linear-algebra operations, Matplotlib (*≥*3.4) [8] for 2D rendering, and BioPython (*≥*1.79) [5] for structure parsing and `NeighborSearch`. RDKit [9] is an optional dependency that, when available, provides chemically accurate 2D ligand coordinates; the package operates fully without it using the PCA projection fallback.

### 2.2 Structure Parsing and Ligand Identification

The `MultiFormatParser` class dispatches to appropriate BioPython parsers (`PDBParser` for `.pdb` and `.pdbqt`; `MMCIFParser` for `.cif` and `.mmcif`) based on file extension. PDBQT files from AutoDock Vina are pre-processed by stripping docking-specific `REMARK` records and normalising atom-name columns to PDB conventions before parsing. Ligand identification follows a hierarchy: (1) the user may supply an explicit residue name via the `–ligand` flag; (2) absent explicit specification, `HybridProtLigMapper` iterates over all hetero-atoms (`HETATM` records), excludes crystallographic water molecules (HOH/WAT) and common buffer components, and selects the largest remaining ligand by heavy-atom count. This heuristic reliably identifies drug-like molecules in standard co-crystal structures.

### 2.3 Interaction Detection

Interaction detection is implemented as a distance-geometry pipeline within the `detect_interactions()` method of `HybridProtLigMapper`. For each of the 15 interaction classes, atom-type selectors and a scientifically validated distance cutoff are defined (Table 1). Candidate interacting atoms are found using BioPython’s `NeighborSearch` spatial index, which partitions protein atoms into a k-d tree and returns all atoms within a specified radius of each ligand atom in *O*(log *n*) time per query. A single search radius equal to the largest applicable cutoff (5.5 Å) is used; individual interaction-type filters are then applied to the returned atom set, reducing combinatorial redundancy.

**Table 1:** Interaction classes detected by PandaMap with validated distance thresholds, atom-type selectors, and primary literature references. Distances are maximum allowed values unless a range is given.

| Interaction Class | Cutoff (Å) | Protein Atoms | Ligand Atoms | Reference |
| --- | --- | --- | --- | --- |
| Hydrogen bond | 2.5–3.5 | N, O (donor/acceptor) | O, N (acceptor/-donor) | PLIP; Auffinger 2004 |
| π–π stacking | 5.5 | Ring C (PHE/-TYR/TRP/HIS) | Ring C (aromatic) | McGaughey 1998 [10] |
| Hydrophobic | 4.0 | C<br>(A/I/L/M/F/P/W/Y/V) | C | Bissantz 2010 [11] |
| Ionic / salt bridge | 5.5 | Charged N (Arg/Lys),<br>O (Asp/Glu) | Opposite charge | Kumar 1999 |
| Cation–π | 5.5 | N (ARG, LYS, HIS) | Aromatic C | Gallivan 1999 [12] |
| Halogen bond | 2.5–3.5 | O, N, S (acceptor) | Cl, Br, I, F | Auffinger 2004 |
| Metal coordination | 2.8 | O, N, S (coordinator) | Metal (Fe, Zn, Mg...) | CSD surveys |
| Carbon–π | 5.5 | Any C | Aromatic ring C | Internal |
| Donor–π | 5.5 | O (ASP, GLU) | Aromatic ring C | Internal |
| Amide–π | 5.5 | N/O (ASN, GLN amide) | Aromatic ring C | Internal |
| Alkyl–π | 5.5 | C (A/V/L/I/M/P) | Aromatic ring C | Internal |
| Attractive charge | 5.5 | Opposite charge pair | Opposite charge pair | Internal |
| Repulsive charge | 4.0 | Same charge pair | Same charge pair | Internal |
| π–cation | 5.5 | Aromatic ring C | Cationic N | Gallivan 1999 [12] |
| Covalent bond | <2.1 | CYS/SER/LYS/HIS re-active atom | Electrophilic atom | Internal |

Ionic/salt-bridge cutoff corrected from 4.0 to 5.5 Å [13]; halogen-bond lower bound corrected from 1.5 to 2.5 Å [14]; H-bond lower bound corrected from 2.4 to 2.5 Å; repulsion cutoff fixed at 4.0 Å (previously ionic × 1.5 = 6.0 Å).

Protein aromatic atoms are identified using the `AROMATIC_RING_ATOMS` dictionary, which stores exact ring-atom name sets for each aromatic residue type: PHE and TYR (CG, CD1, CD2, CE1, CE2, CZ), TRP (CG, CD1, CD2, CE2, CE3, CZ2, CZ3, CH2, NE1), and HIS (CG, ND1, CD2, CE1, NE2). This atom-name filter ensures that *β*-carbons and other non-ring atoms do not generate false *π*-system contacts. Similarly, ligand ring atoms are determined by the topological ring-detection algorithm (Section 2.4) rather than element-type heuristics.

### 2.4 Ligand Ring Detection

Correct identification of ring atoms is prerequisite for *π*-system interaction detection. PandaMap implements a topology-based ring-detection algorithm using iterative leaf-node pruning on the heavy-atom bond graph (Algorithm 1). The algorithm runs in *O*(*n* + *m*) time, where *n* is the number of atoms and *m* the number of bonds, and correctly identifies all ring atoms regardless of ring size or fused-ring topology, without requiring SMILES strings or a chemistry toolkit. Only atoms whose topological degree remains *≥*2 after complete pruning are classified as ring members, a necessary and sufficient condition for cycle membership.

**Listing 1:**
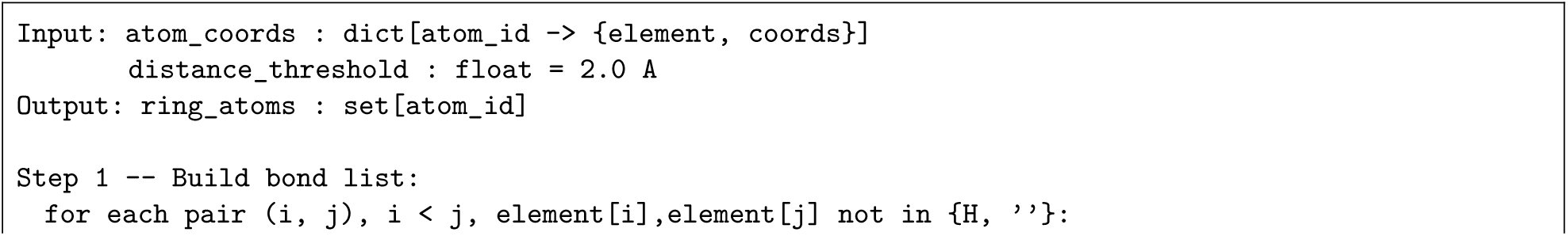
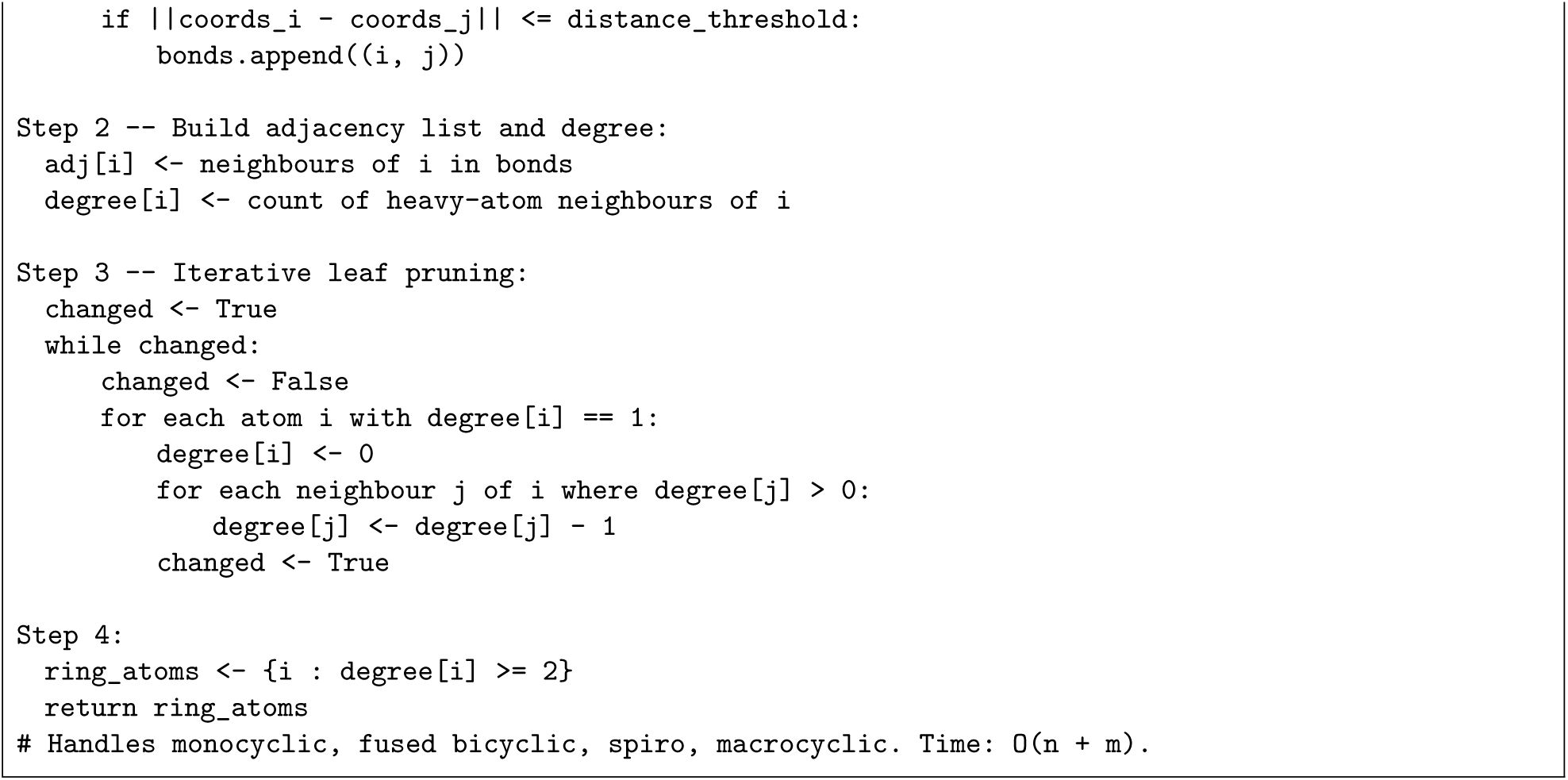
Algorithm 1 – Ligand ring detection (leaf-node pruning).

### 2.5 2D Ligand Coordinate Generation

Two-dimensional ligand coordinates for the interaction diagram are generated via a two-tier strategy. When RDKit is installed, an `RWMol` object is constructed from the heavy-atom bond topology and `AllChem.Compute2DCoords()` is invoked to produce chemically accurate 2D coordinates with ring planarity, stereochemistry awareness, and collision resolution (Algorithm 2, Tier 1). When RDKit is unavailable, PandaMap falls back to a PCA-based 3D*→*2D projection that operates on NumPy alone (Algorithm 2, Tier 2). The detected interactions are identical either way, since detection works from the 3D coordinates and never uses the 2D layout; what differs is the depiction. The PCA projection flattens a 3D conformer onto its two principal axes, so rings are not drawn with regular geometry and atoms can overlap where the molecule folds back on itself. It keeps the package usable with no chemistry toolkit installed, but RDKit is worth having if the diagram is destined for a figure. Figure S1 compares the two on PDB 1US0.

**Listing 2:**
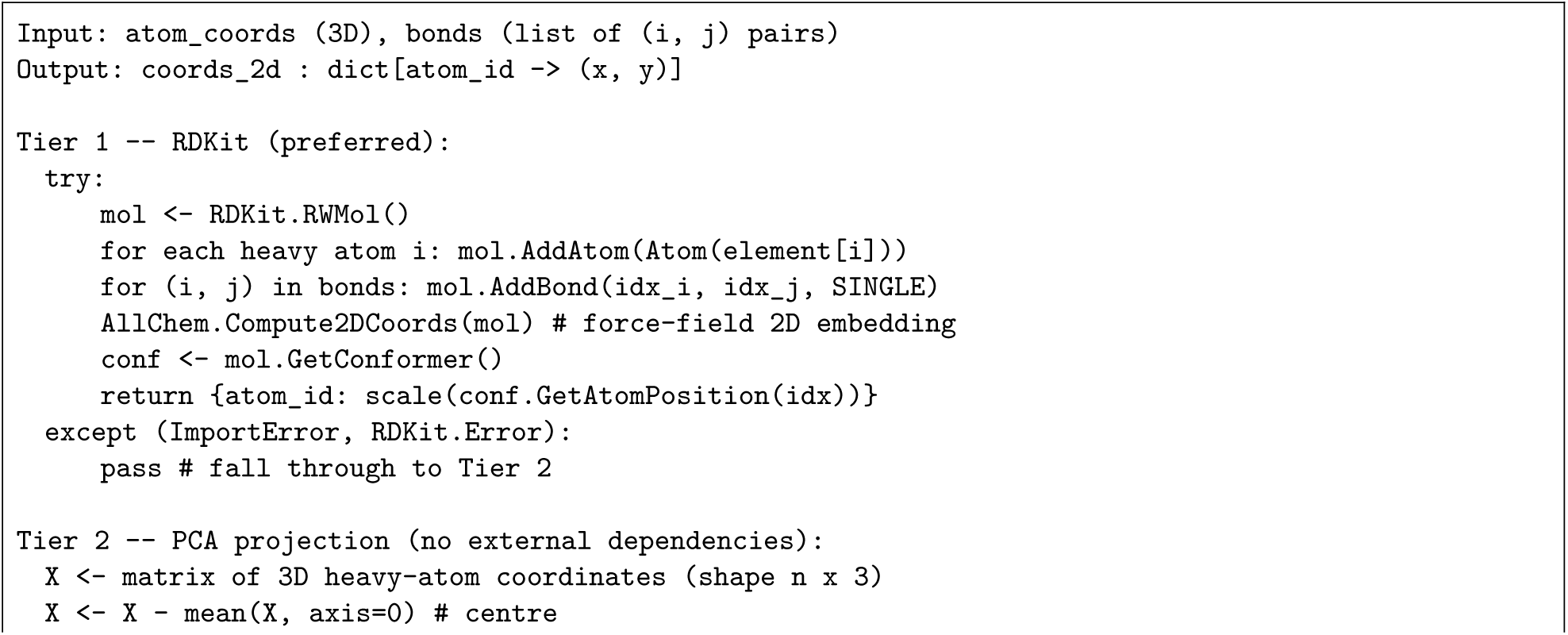
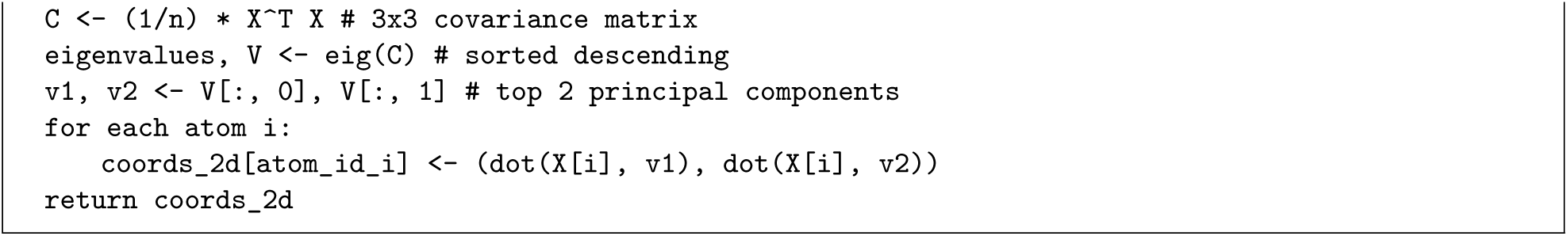
Algorithm 2 – 2D coordinate generation (RDKit → PCA fallback).

### 2.6 Hydrogen-Bond Geometry

A hydrogen bond is not defined by donor–acceptor distance alone; the D–H*· · ·* A angle matters, and a contact that is 3.0 Å but bent is not the same thing as a contact that is 3.0 Å and linear. PLIP enforces a donor-angle minimum of 100*^◦^*, which it can do because it protonates every structure with OpenBabel before profiling.

PandaMap takes a narrower position, for a reason worth stating plainly. When the deposited coordinates already contain hydrogens (neutron structures, ultrahigh-resolution X-ray, NMR ensembles, or models a user has protonated themselves), the true D–H*· · ·* A angle is measured for both candidate donor directions, the more favourable is kept, and the contact is rejected below 100*^◦^*. This is the same criterion PLIP applies, on the same evidence.

When hydrogens are absent, as in most X-ray depositions, PandaMap falls back to the distance window alone. We implemented a heavy-atom proxy that infers an approximate H direction from the donor’s covalent neighbours, and it is available behind a flag, but it is off by default because it fails in a specific and instructive way: without hydrogens it cannot tell which partner is the donor, so a genuine bond to a carbonyl acceptor is scored as though the carbonyl were donating. Applied as a hard filter to PDB 1ELS it removes nine of ten hydrogen bonds, all of which PLIP confirms. A criterion that discards correct answers is worse than no criterion, and adding hydrogens is the honest fix; we would rather users protonate their structures than trust a proxy that looks rigorous and is not.

Every reported hydrogen bond therefore carries both the measured angle and its provenance (`explicit_H` or `heavy_atom`), so downstream analysis can tell which rule applied. On the 20-model NMR ensemble of PDB 1ACA, where hydrogens are present, all six retained contacts lie between 128*^◦^* and 173*^◦^*.

### 2.7 Empirical Binding-Affinity Estimation

PandaMap provides an empirical binding free-energy estimate (Δ*G*, kcal/mol) using a count-based scoring function adapted from the Böhm [15] and X-Score [16] frameworks. Three improvements over a naïve atom-count sum are incorporated. First, per-residue deduplication is applied: for each interaction type, only the closest atom-level contact to each protein residue is retained, so that multi-atom residues (e.g. a phenylalanine ring with six carbons all within alkyl–*π* range) contribute at most one contact per class. Second, a linear distance-decay function scales each contribution between full weight at the ideal contact distance and zero weight at the detection cutoff, so that contacts at the boundary of the detection radius do not receive the same weight as optimal-geometry contacts. Third, a rotatable-bond penalty of +0.5 kcal/mol per freely rotatable non-ring ligand bond reflects the conformational entropy cost of restricting flexible ligands on binding [15]. The full scoring equation is

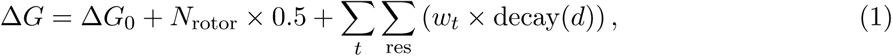

where Δ*G*_0_ = +2.9 kcal/mol is the base desolvation and translational/rotational entropy constant [15]; *N*_rotor_ is the number of freely rotatable ligand bonds; and the double sum runs over interaction types *t* and unique residues. The decay function is decay(*d*) = 1 for *d ≤ d*_ideal_, and max(0, (*d*_max_ *− d*)*/*(*d*_max_ *− d*_ideal_)) for *d > d*_ideal_. Qualitative *K_d_* labels are assigned using Δ*G* = *−RT* ln *K_d_* at 298 K (*RT* = 0.593 kcal/mol): Δ*G < −*12 (*K_d_* ≲ nM); *< −*9 (nM–*µ*M); *< −*6 (*µ*M); *< −*3 (mM); *≥ −*3 (very weak). Systematic errors of *±*2–3 kcal/mol are expected; the estimate is intended for rapid relative ranking, not absolute *K_d_* prediction (see Section 4.3).

### 2.8 Multi-Frame Trajectory Analysis

The module-level `analyze_trajectory` function processes multi-model PDB files produced by MD simulation or NMR ensemble refinement. Each MODEL record is extracted using BioPython’s PDBIO and a single-model selector, written to a temporary file, and analysed independently by `HybridProtLigMapper`. Per-frame interaction counts and Δ*G* estimates are collected. At the end of the trajectory, per-residue interaction occupancies (fraction of frames in which each contact is observed) and mean interaction counts are computed and exported to a CSV file. Optionally, a 2D interaction diagram is generated for each frame.

### 2.9 Solvent Accessibility and Visualization

Residues bordering the solvent-exposed face of the binding pocket are marked with a cyan halo in the 2D diagram. PandaMap implements a layered SASA estimation strategy: when the DSSP binary [17] is available, BioPython’s DSSP interface is invoked and relative accessible surface-area values are read directly; otherwise a geometric estimator based on atom density around each residue C*α* provides a dependency-free fallback. (An internal Shrake–Rupley implementation [18] using Fibonacci-distributed sphere points is also included in the package.) The 2D interaction diagram is rendered as a Matplotlib figure with protein residues arranged in a circular layout around the central 2D ligand depiction. Interaction arcs are drawn as `FancyArrowPatch` curves; multiple interactions to the same residue are fanned out using an alternating-sign curvature offset to prevent marker overlap. The 3D HTML output is a self-contained file embedding the structure as a base64-encoded PDB string within a 3Dmol.js viewer (Figure 2).

**Figure 2:**
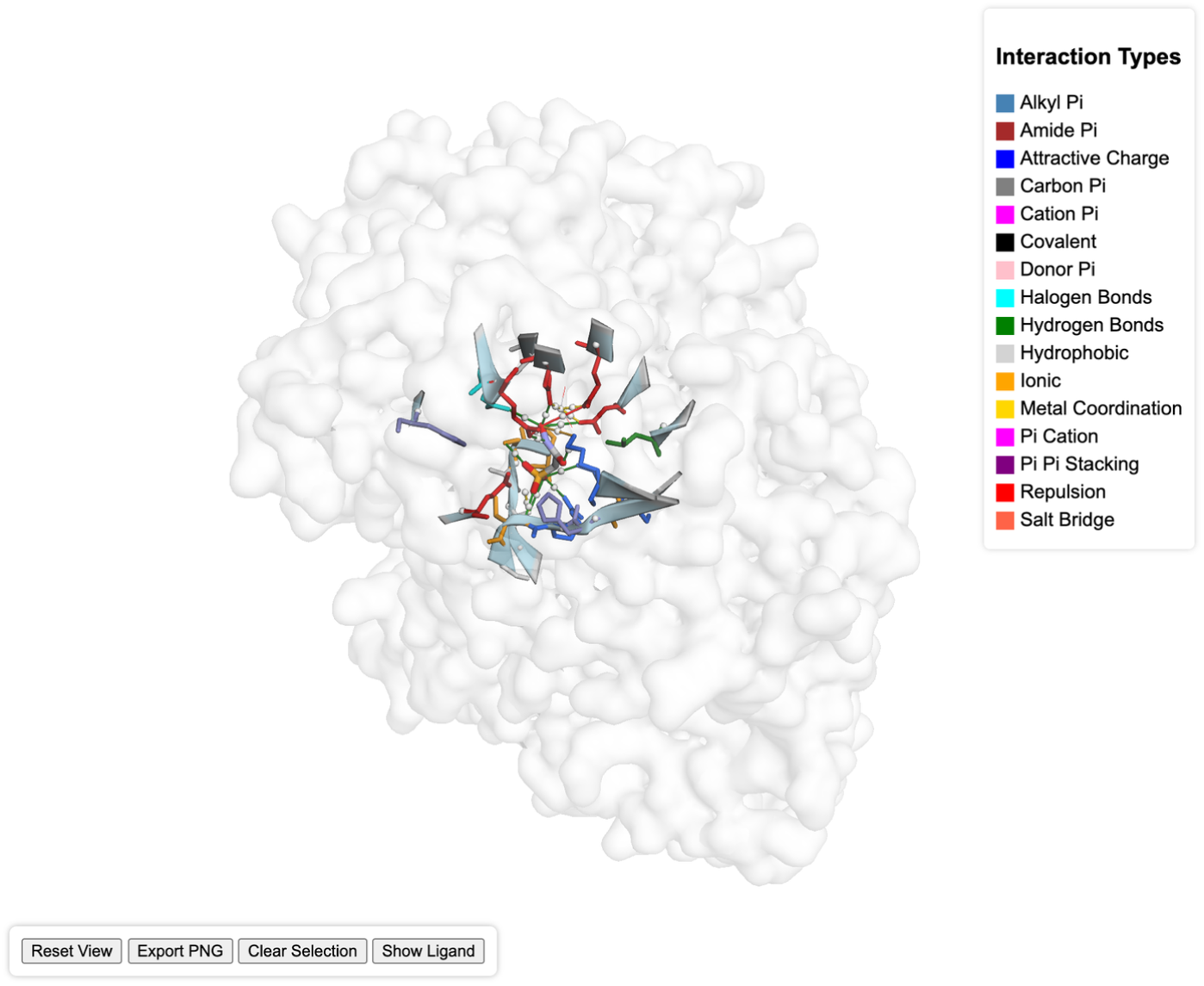
Self-contained interactive 3D viewer generated by the --3d flag for the enolase–PAH complex (PDB 1ELS), rendered with 3Dmol.js. The protein is shown as a translucent surface, the ligand as sticks, and detected interactions as colour-coded dashed lines keyed to the legend; the page provides interactive controls (rotate, reset, export, toggle ligand). The figure is a static capture of the otherwise fully interactive HTML output included as Supplementary File S1.

## 3 Results and Validation

PandaMap was validated on three structurally diverse, well-characterised complexes spanning metalloenzyme, protein-kinase, and oxidoreductase chemistry. All results below were produced with PandaMap 4.3.0 using the commands listed in the Supplementary Information; reported distances and counts are taken verbatim from the generated text reports, and the full interaction counts for all three complexes are collected in Table 2.

**Table 2:** PandaMap interaction summary for the three benchmark complexes. Values are filtered interaction counts after per-residue deduplication; “–” denotes not detected or not applicable. Δ*G* estimated using the empirical scoring function (Section 2.7) at 298 K.

| Interaction class | 1ELS (PAH) | 1M17 (erlotinib) | 1US0 (IDD594) |
| --- | --- | --- | --- |
| Hydrogen bonds | 10 | 4 | 3 |
| π-π stacking | – | – | 9 |
| Hydrophobic | – | 6 | 7 |
| Metal coordination | 4 | – | – |
| Ionic / salt bridge | 2 | 1 | 2 |
| Halogen bond | – | – | 4 |
| π-cation | – | 1 | – |
| Carbon-π | 1 | – | 10 |
| Donor-π | – | 2 | – |
| Amide-π | – | 1 | – |
| Alkyl-π | 1 | 8 | 8 |
| Attractive charge | 2 | 1 | 2 |
| $\Delta G$ estimate (kcal/mol) | –18.10 | –0.86 | –13.92 |
| Interacting residues | 12 | 14 | 15 |

### 3.1 Case Study 1: Enolase Transition-State Complex (PDB 1ELS)

PDB 1ELS is yeast enolase bound to phosphonoacetohydroxamate (ligand PAH), a transition-state analogue, together with the two catalytic Mn^2+^ ions [19]. It is a useful stress test because almost everything of interest is electrostatic: a phosphonate group buried in a cluster of acidic residues, bridged by metal.

PandaMap found 12 interacting residues (Figure 3a). Ten hydrogen bonds span 2.60–3.46 Å, among them GLU168 at 2.66 Å, SER375 at 2.82 Å and ARG374 at 2.98 Å. Four metal-coordination contacts at 2.12–2.24 Å pick out GLY37, ASP246, GLU295 and ASP320; these are the residues Zhang and co-workers identify as the divalent-metal site, and the same set across the enolase superfamily. Two charge–charge contacts anchor the phosphonate to LYS345 and ARG374.

**Figure 3:**
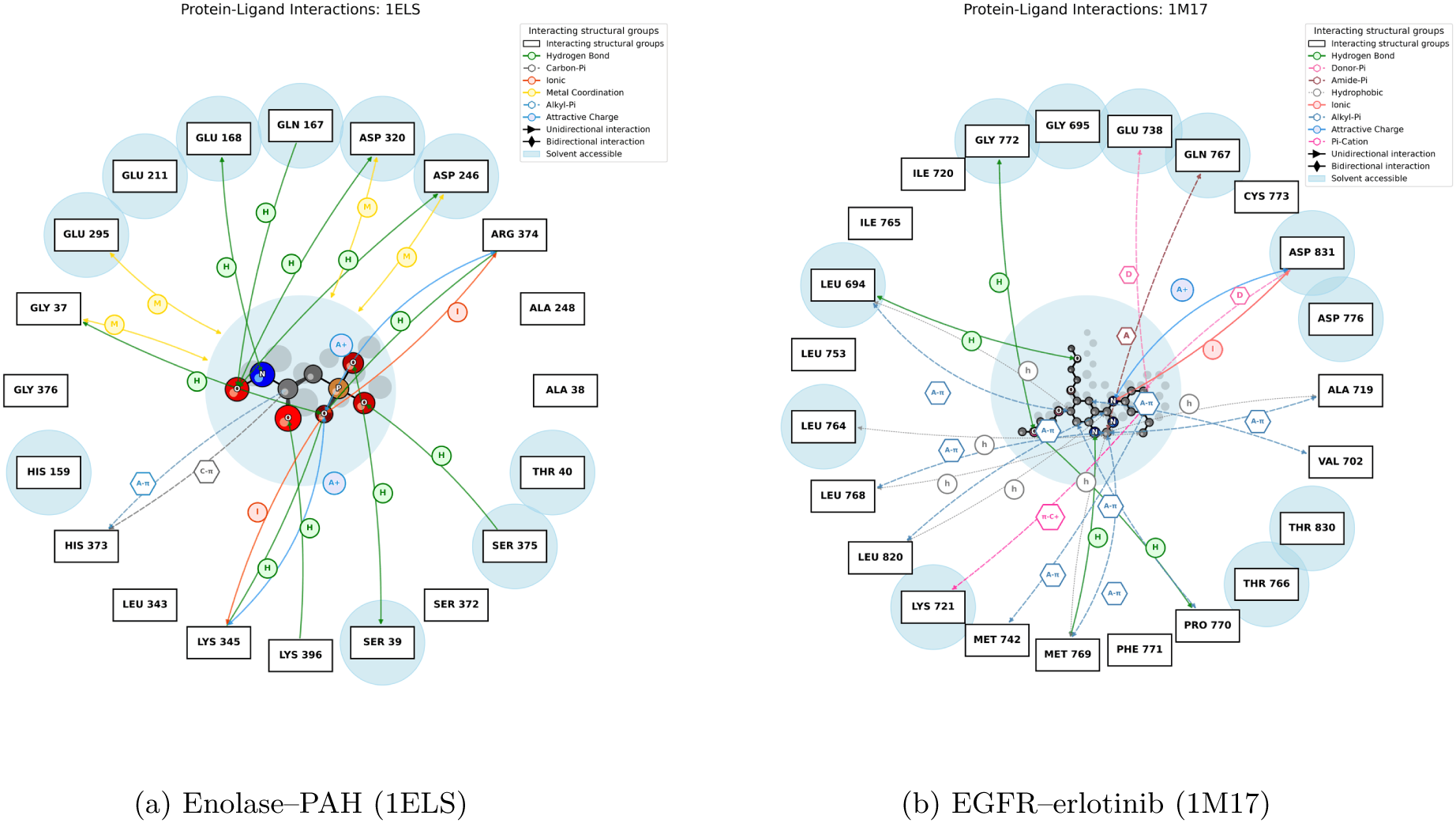
Interaction diagrams for the enolase and EGFR benchmarks, discussed in Sections 3.1 and 3.2. (a) The four Mn^2+^-coordinating residues (GLY37, ASP246, GLU295, ASP320) and the phosphonate charge network. (b) The EGFR hinge hydrogen bond to MET769 at 2.70 Å with the surrounding hydrophobic and alkyl–*π* contacts of the adenine pocket. The aldose reductase diagram is Figure 5.

The empirical estimate is Δ*G ≈ −*18.1 kcal/mol. Transition-state analogues bind tightly, so the direction is right, though the magnitude should not be read too literally (Section 4.3).

### 3.2 Case Study 2: EGFR Kinase–Erlotinib Complex (PDB 1M17)

PDB 1M17 is the EGFR tyrosine-kinase domain with erlotinib bound (ligand AQ4) [20]. Erlotinib’s binding mode is about as well characterised as a kinase inhibitor’s can be, which makes it a good test of whether a tool recovers the contacts a medicinal chemist would draw by hand.

It does. All 14 interacting residues are ATP-pocket residues, and the diagnostic hinge hydrogen bond from the quinazoline N1 to the MET769 backbone appears at 2.70 Å (Figure 3b), with three further hydrogen bonds to LEU694 (3.27 Å), PRO770 (3.21 Å) and GLY772 (3.24 Å). Six hydrophobic and eight alkyl–*π* contacts line the adenine pocket, and a *π*–cation contact reaches the conserved *β*3 lysine, LYS721, at 3.70 Å.

The scoring function is another matter. It returns Δ*G ≈ −*0.86 kcal/mol for a low-nanomolar inhibitor. We report this rather than quietly dropping the case: a +3.0 kcal/mol rotatable-bond penalty from erlotinib’s two methoxyethoxy chains dominates the sum, and nothing in a count-based function knows that the hinge hydrogen bond is worth more than its distance suggests. The contact topology is right; the number is not. Section 4.3 takes this up.

### 3.3 Case Study 3: Aldose Reductase–IDD594 Complex (PDB 1US0)

PDB 1US0 is human aldose reductase with the inhibitor IDD594 (ligand LDT), determined at 0.66 Å [21]. The structure is the standard reference for protein–ligand halogen bonding, because the inhibitor’s selectivity is usually attributed to a short bromine*· · ·* oxygen contact at THR113.

PandaMap recovers that contact at 2.97 Å, the same distance PLIP reports, along with three C– F*· · ·* O contacts to VAL47 (3.01 Å), ALA299 (3.26 Å) and LEU300 (3.29 Å). Three hydrogen bonds reach the catalytic anion-binding site (HIS110 2.66 Å, TYR48 2.73 Å, TRP111 3.06 Å), and the carboxylate head group makes charge–charge contacts with HIS110 and LYS77. Nine *π*–*π* stacking, ten carbon–*π*, eight alkyl–*π* and seven hydrophobic contacts fill out an unusually aromatic pocket lined by four tryptophans (Figure 5, Figure 4). The estimate is Δ*G ≈ −*13.9 kcal/mol, which is at least in the right range for a low-nanomolar inhibitor.

**Figure 4:**
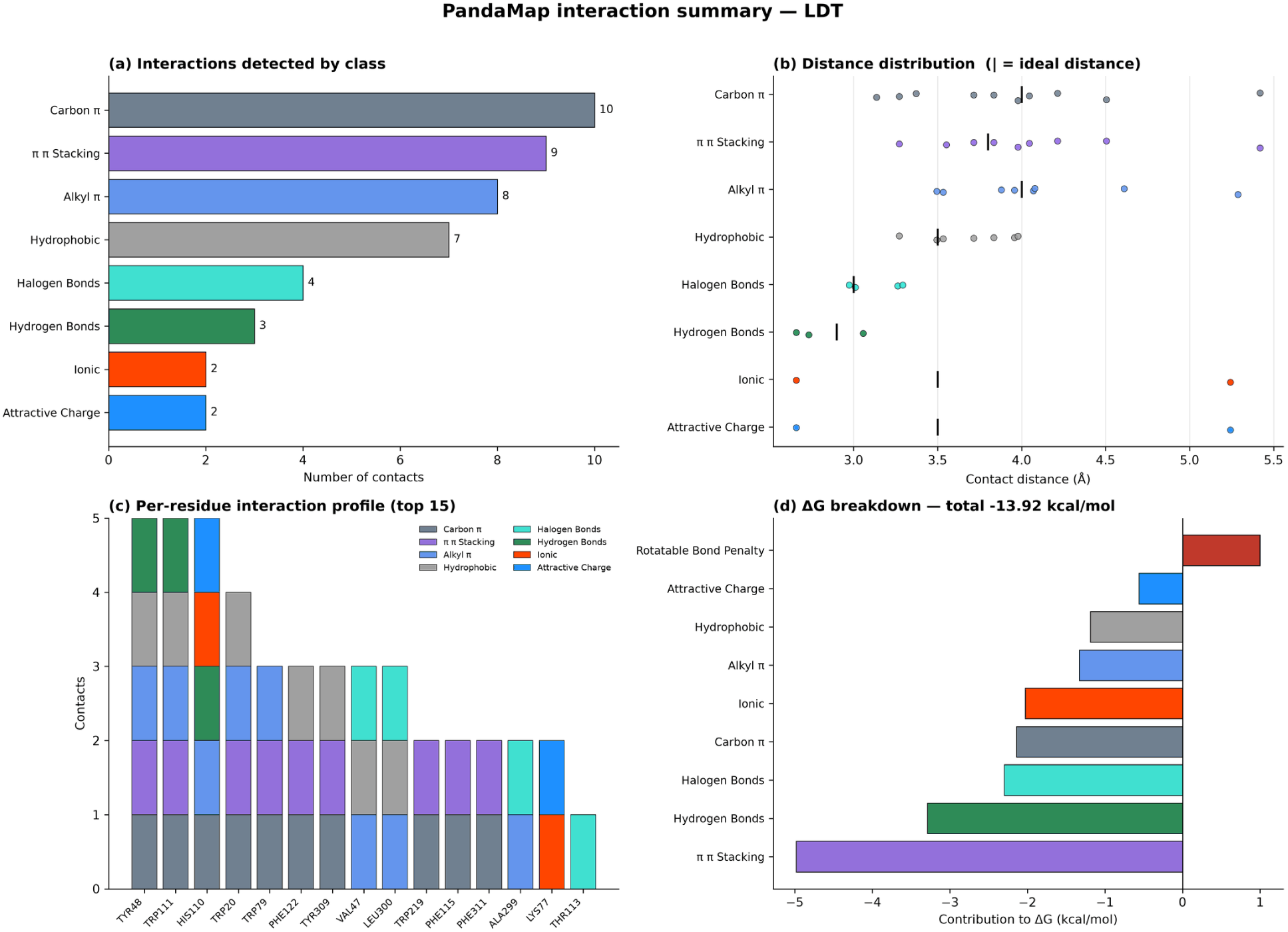
Graphical interaction report for PDB 1US0, produced by –plots alongside the diagram and the text report. (a) Contacts per class. (b) Distance distribution per class, with the ideal contact distance marked; contacts sitting near the detection cutoff are distinguishable at a glance from those with near-optimal geometry. (c) Per-residue profile, stacked by class; HIS110 and TYR48 stand out as the hotspot residues. (d) Per-class contribution to the empirical Δ*G*, with penalties in red. Panels (b) and (d) are the ones users have found most informative, since neither is recoverable from the 2D diagram.

**Figure 5:**
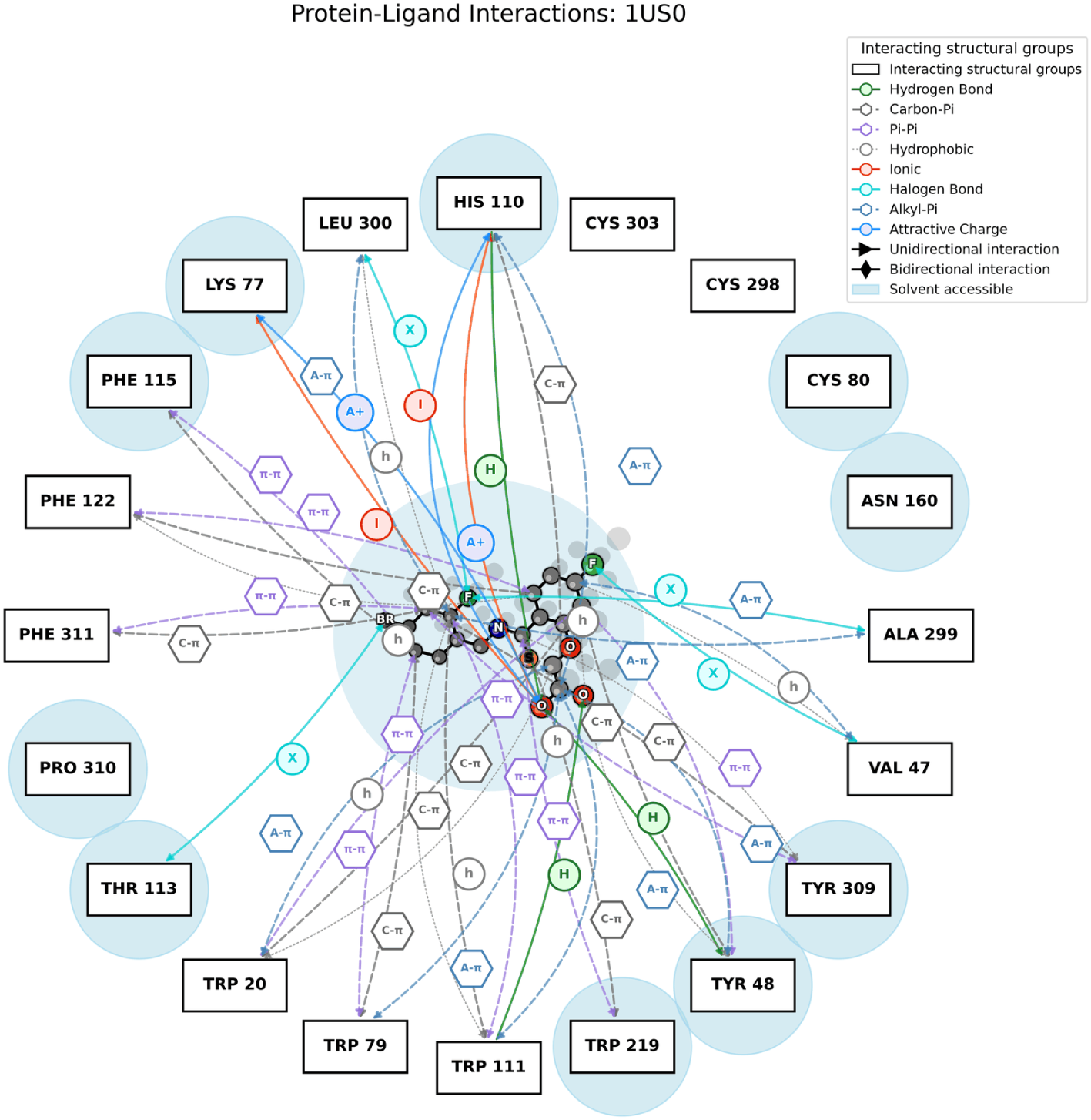
PandaMap 2D interaction diagram for aldose reductase–IDD594 (PDB 1US0). Protein residues sit radially around the ligand depiction and arcs are coloured by interaction class. The bromine*· · ·* THR113 halogen bond and the three C–F*· · ·* O contacts are drawn in cyan (**X**); hydrogen bonds to the catalytic anion-binding site (TYR48, HIS110, TRP111) in green. Cyan haloes mark solvent-accessible residues. Ligand coordinates come from RDKit; compare Figure S1 for the dependency-free fallback.

Two of these findings were invisible in release 4.2.1 and are worth flagging, because they were not detected by release 4.2.x (Section 5): the bromine contact was missed entirely, and so were both carboxylate salt bridges.

### 3.4 Comparison with PLIP

We profiled all three complexes independently with PLIP 2.3.0 [2, 3] and compared the contacts residue by residue (Table 3).

**Table 3:**
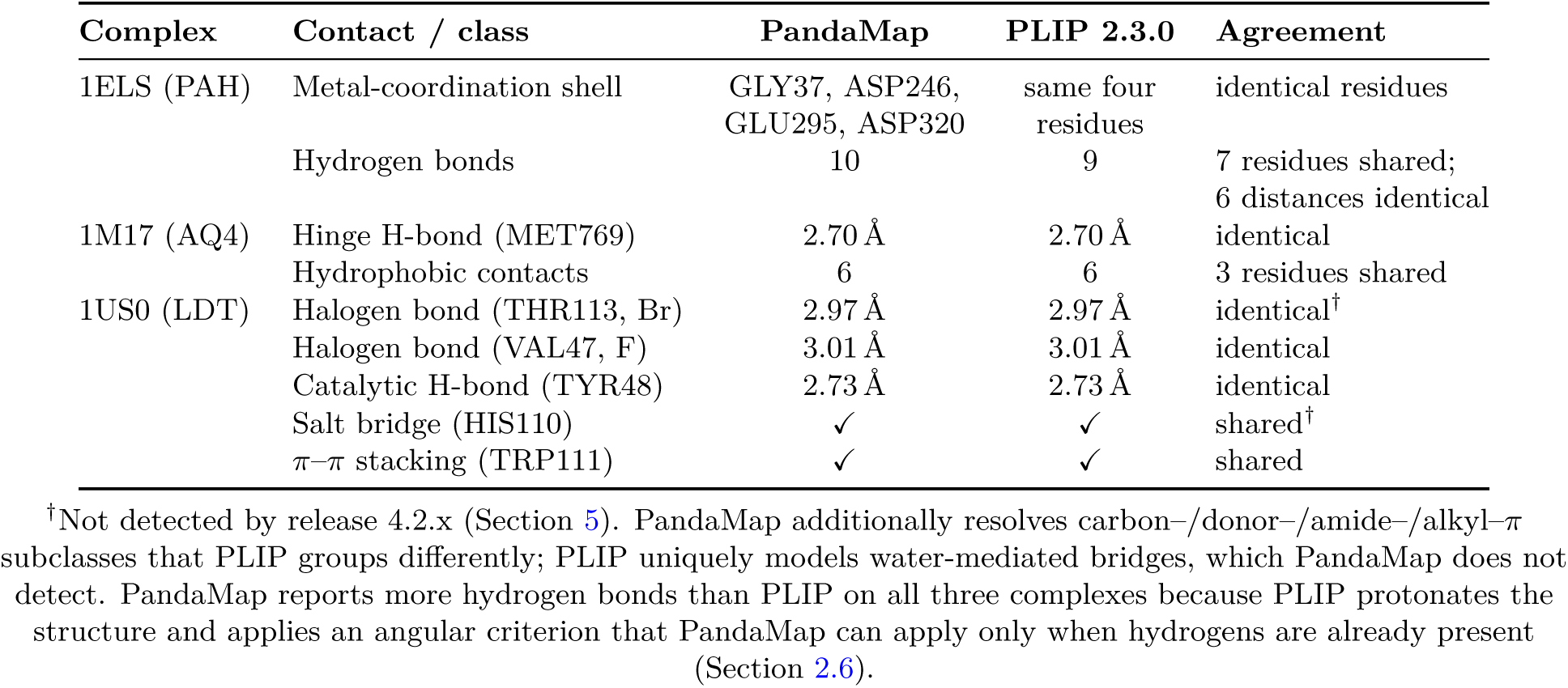
Direct comparison of diagnostic contacts detected by PandaMap and PLIP (v2.3.0) for the three benchmark complexes. “Shared” counts residues detected by both tools in the same interaction class.

| Complex | Contact / class | PandaMap | PLIP 2.3.0 | Agreement |
| --- | --- | --- | --- | --- |
| 1ELS (PAH) | Metal-coordination shell | GLY37, ASP246, GLU295, ASP320 | same four residues | identical residues |
|  | Hydrogen bonds | 10 | 9 | 7 residues shared; 6 distances identical |
| 1M17 (AQ4) | Hinge H-bond (MET769) | 2.70 Å | 2.70 Å | identical |
|  | Hydrophobic contacts | 6 | 6 | 3 residues shared |
| 1US0 (LDT) | Halogen bond (THR113, Br) | 2.97 Å | 2.97 Å | identical<sup>†</sup> |
|  | Halogen bond (VAL47, F) | 3.01 Å | 3.01 Å | identical |
|  | Catalytic H-bond (TYR48) | 2.73 Å | 2.73 Å | identical |
|  | Salt bridge (HIS110) | ✓ | ✓ | shared<sup>†</sup> |
| | π-π stacking (TRP111) | ✓ | ✓ | shared |

<sup>†</sup>Not detected by release 4.2.x (Section 5). PandaMap additionally resolves carbon-/donor-/amide-/alkyl- $\pi$ subclasses that PLIP groups differently; PLIP uniquely models water-mediated bridges, which PandaMap does not detect. PandaMap reports more hydrogen bonds than PLIP on all three complexes because PLIP protonates the structure and applies an angular criterion that PandaMap can apply only when hydrogens are already present (Section 2.6).

Where the two tools assign the same class to the same pair, the distances agree to the hundredth of an Ångström, which is what one would hope for given that both measure the same interatomic separations. The EGFR hinge bond to MET769 is 2.70 Å in both. For IDD594, PLIP reports two halogen bonds, VAL47 at 3.01 Å and THR113 at 2.97 Å, and PandaMap now reproduces both exactly, along with two further C–F*· · ·* O contacts to ALA299 and LEU300 that fall inside its cutoff. For enolase, six of PandaMap’s ten hydrogen bonds match PLIP distances exactly (GLU168 2.66, SER375 2.82, ARG374 2.98, SER39 3.06, GLN167 3.10, LYS345 3.21 Å), and both tools independently identify the same four metal-coordinating residues: GLY37, ASP246, GLU295 and ASP320.

The counts differ, and it is worth being precise about why rather than presenting the overlap alone. PandaMap reports more hydrogen bonds than PLIP on all three complexes: ten versus nine on 1ELS, four versus one on 1M17, three versus one on 1US0. The gap is largest where PLIP applies angular criteria that PandaMap cannot: PLIP protonates the structure with OpenBabel before profiling and then enforces a donor-angle cutoff, whereas PandaMap measures the true D–H*· · ·* A angle only when the deposited coordinates already contain hydrogens (Section 2.6). None of the three X-ray benchmarks does. On 1M17 and 1US0, PandaMap is therefore applying a distance criterion where PLIP applies distance and geometry, and the extra contacts it reports should be read as candidates rather than as disagreements about the same evidence.

In the other direction, PLIP models water-mediated bridges, which PandaMap does not detect at all, and PandaMap resolves carbon–*π*, donor–*π*, amide–*π* and alkyl–*π* subclasses that PLIP groups differently. The two vocabularies are not in conflict; they were built for different purposes.

### 3.5 Multi-Frame Trajectory Analysis

To exercise the trajectory module on real multi-model data, the 20-model solution-NMR ensemble of acyl-coenzyme A binding protein (ACBP) bound to palmitoyl-coenzyme A (PDB 1ACA, ligand COA) [22] was processed with `analyze_trajectory`. PandaMap analysed all 20 models independently in a single call and exported a per-frame summary CSV and per-residue occupancy statistics (Figure 6). The per-frame empirical estimate was Δ*G* = *−*8.96 *±* 2.04 kcal/mol across the ensemble (range *−*12.2 to *−*4.4; mean 6.4 hydrogen bonds per frame). The occupancy analysis separates a core of contacts present in at least 80% of models (ALA9, VAL12, LYS13, ILE27, TYR28, TYR31, LYS32, LYS54, TYR73) from ones that appear only intermittently (LEU15 0.30, MET24 0.20, LYS16 0.05).

**Figure 6:**
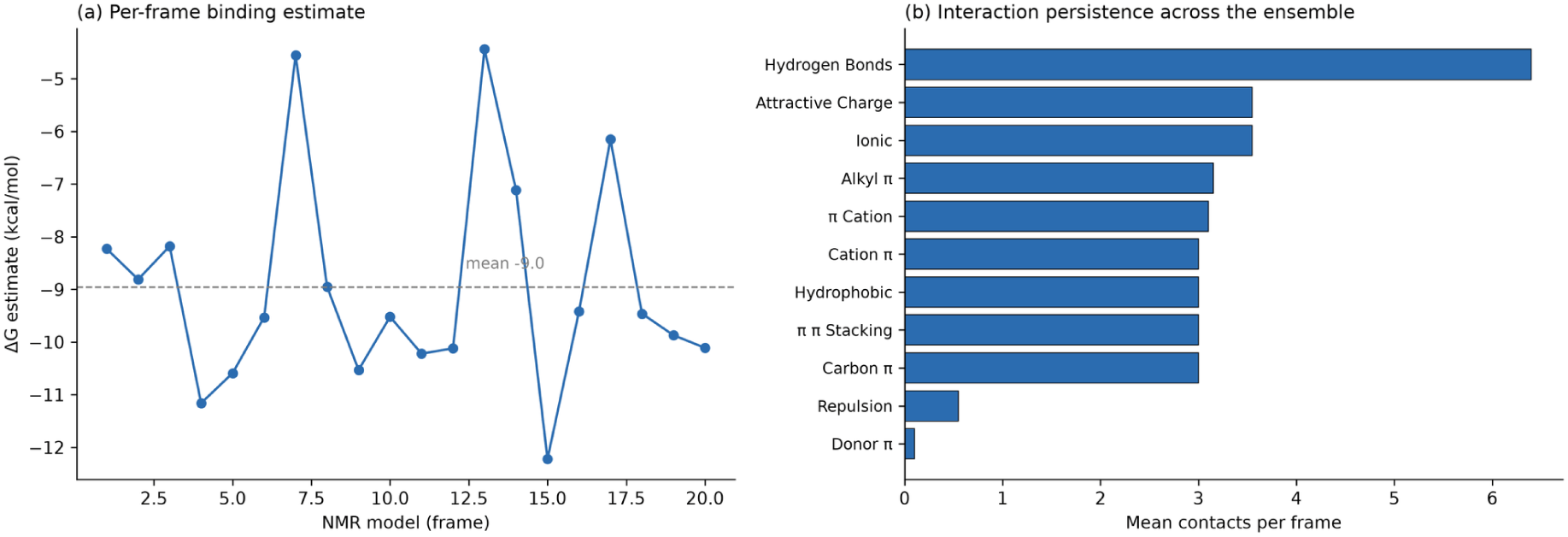
Multi-frame trajectory analysis of the 20-model NMR ensemble of acyl-coenzyme A binding protein–palmitoyl-coenzyme A (PDB 1ACA) [22]. (a) Per-frame empirical Δ*G* estimate across the 20 models (dashed line: mean *−*8.96 kcal/mol). (b) Per-residue contact occupancy (fraction of frames in which each residue contacts the ligand), separating stable (*≥*0.8), intermittent, and transient (*<*0.4) contacts. Both outputs are produced automatically and written to a summary CSV.

The stable set is a useful check on the method, because it was not chosen by us: Kragelund and co-workers identify Ala9, Tyr28, Lys32 and Lys54 as the residues hydrogen bonding or salt bridging to the adenosine-3*^′^*-phosphate of the bound ligand [22], and all four fall in the persistent group. Separating persistent from transient contacts is the main reason to run the occupancy analysis on an NMR ensemble or an MD trajectory; a single frame cannot tell you which contacts are structural and which are incidental.

## 4 Discussion

PandaMap fills a well-defined niche in the structural-bioinformatics software ecosystem: rapid, zero-configuration, publication-ready interaction mapping from within a standard Python environment. The design philosophy prioritises (i) minimal mandatory dependencies, (ii) correctness of interaction detection over exhaustive coverage, and (iii) visual clarity of output.

### 4.1 Distance-Cutoff Corrections

The original implementation used a 2.4 Å lower bound for hydrogen bonds, a 4.0 Å cutoff for ionic interactions, and a 1.5 Å lower bound for halogen bonds. The corrected values (2.5, 5.5, and 2.5 Å respectively) are grounded in published crystallographic surveys and align with PLIP’s thresholds. The 2.4 Å lower bound was below the van der Waals contact minimum for N*· · ·* O pairs and generated spurious contacts in strained geometries. The 4.0 Å ionic cutoff missed a substantial fraction of genuine salt bridges identified by Kumar & Nussinov [13] at 4.5–5.5 Å. The 1.5 Å halogen lower bound was within the covalent-bond range and produced false positives for every halogenated ligand with a protein contact.

### 4.2 Ring Detection and Aromatic-Atom Filtering

The leaf-node-pruning ring-detection algorithm replaces the previous element-type heuristic, which classified any ligand carbon as aromatic if it belonged to a residue type known to be aromatic. This caused false *π*-system contacts for non-ring carbons in partially saturated or heteroaromatic ligands. The topology-based approach is exact for any bond topology without requiring SMILES. On the protein side, the `AROMATIC_RING_ATOMS` atom-name filter eliminates false contacts from *β*-carbons, which are within 5.5 Å of ligands in many structures but are not part of the aromatic *π* system.

### 4.3 Empirical Δ*G* Estimation

Three features of the scoring function are defensible. Per-residue deduplication stops a phenylalanine ring from contributing six times what a single contact should contribute. Distance decay reflects the fact that a contact at the edge of the detection radius is not worth the same as one at ideal geometry. The rotatable-bond penalty is a crude nod to the conformational entropy a flexible ligand gives up on binding.

None of that makes the output a binding affinity, and the EGFR benchmark shows why. Erlotinib inhibits EGFR at low nanomolar concentrations. PandaMap scores it at *−*0.86 kcal/mol, which would be a non-binder. The arithmetic is easy to follow: two methoxyethoxy chains contribute six rotatable bonds and a +3.0 kcal/mol penalty, while the hinge hydrogen bond that actually determines the binding mode is worth *−*1.2 kcal/mol, the same as any other hydrogen bond in the table. A count-based function has no way to know that one hydrogen bond is load-bearing and another is incidental.

The failure is systematic rather than a quirk of this ligand. Weights taken from the literature rather than fitted to data will misrank any comparison across chemotypes, and the *±*2–3 kcal/mol figure we quote is a floor, not an error bar. Within a congeneric series of similar flexibility the ordering is more likely to be meaningful. Beyond that, use MM-GB/PBSA or free-energy perturbation and treat this number as a sanity check.

### 4.4 Remaining Limitations and Future Work

Four limitations are worth stating plainly, since users will meet all of them.

The angular hydrogen-bond criterion applies only where hydrogens exist in the deposited coordinates (Section 2.6). For the majority of X-ray structures PandaMap is a distance filter, and its hydrogen-bond counts will exceed PLIP’s for that reason alone. Users who care about geometric fidelity should protonate their structures first; adding an OpenBabel-based protonation step, as PLIP does, is the obvious remedy and is planned.

*π*–*π* stacking uses atom-to-atom distance rather than ring-centroid separation with a dihedral check, so face-to-face, T-shaped and parallel-displaced geometries are not distinguished and the counts are inflated relative to a centroid-based definition. The ten carbon–*π* and nine *π*–*π* contacts reported for 1US0 should be read as a description of an aromatic environment, not as ten and nine discrete stacking events.

Metal coordination uses one 2.8 Å cutoff for every metal, where element-specific ranges (Zn–N 2.0–2.2 Å; Ca–O 2.3–2.9 Å) would be more discriminating.

Finally, and most importantly for anyone tempted to use the Δ*G* output quantitatively: the weights are not regression-fitted to any benchmark. They are literature-derived per-contact estimates, and the erlotinib case in Section 4.3 shows what that costs. Counting each electrostatic contact once, as 4.3.0 does, makes the function internally consistent; it does not make it accurate. Treat the number as a coarse ranking heuristic and use MM-GB/PBSA or free-energy perturbation when the value matters.

## 5 Conclusions

PandaMap produces a publication-quality protein–ligand interaction diagram, an interactive 3D view, a text report, a machine-readable CSV, and a graphical summary from one command, with four dependencies and no configuration. On three benchmark complexes spanning metalloenzyme, kinase and oxidoreductase chemistry it recovers the contacts each structure is known for: the enolase metal-coordination shell, the EGFR hinge hydrogen bond, and the IDD594 bromine halogen bond, the latter two at distances identical to those reported by PLIP.

Two caveats bear repeating, because they bound what the tool should be used for. The angular hydrogen-bond criterion is exact only when the structure carries explicit hydrogens; for most X-ray depositions PandaMap applies a distance filter, and users who need geometric fidelity should protonate first. The empirical Δ*G* is a ranking heuristic built from literature weights, not a fitted affinity model, and the EGFR case shows how far it can miss on a flexible ligand.

Within those limits, the package covers the case most structural and medicinal chemists actually have: one structure, one ligand, and the need for a figure and a contact list without building a pipeline first. PandaMap 4.3.0 is available on PyPI under the MIT licence.

## Acknowledgements

The author thanks the developers of BioPython, Matplotlib, RDKit, and 3Dmol.js, whose libraries form the foundation of PandaMap. Benchmark PDB structures were obtained from the RCSB Protein Data Bank (rcsb.org). This work received no external funding.

## Code and Data Availability

PandaMap is free and open-source software released under the MIT licence. The released package is distributed via PyPI (pip `install pandamap`==4.3.0); source code, issue tracker, and development history are hosted at https://github.com/pritampanda15/PandaMap. The version documented in this manuscript is v4.3.0. The benchmark structures 1ELS, 1M17, 1US0, and 1ACA are available from the RCSB Protein Data Bank (https://www.rcsb.org). The exact commands used to generate every result and figure in this manuscript are listed in the Supplementary Information, and the resulting interaction reports are included in the repository.

## Author Contributions

P.K.P. conceived the software, wrote the code, performed all analyses, and wrote the manuscript.

## Ethics

This study used only publicly available structural data from the RCSB Protein Data Bank and involved no human participants, human data, or animal subjects.

## Supplementary Information

**Figure S1:**
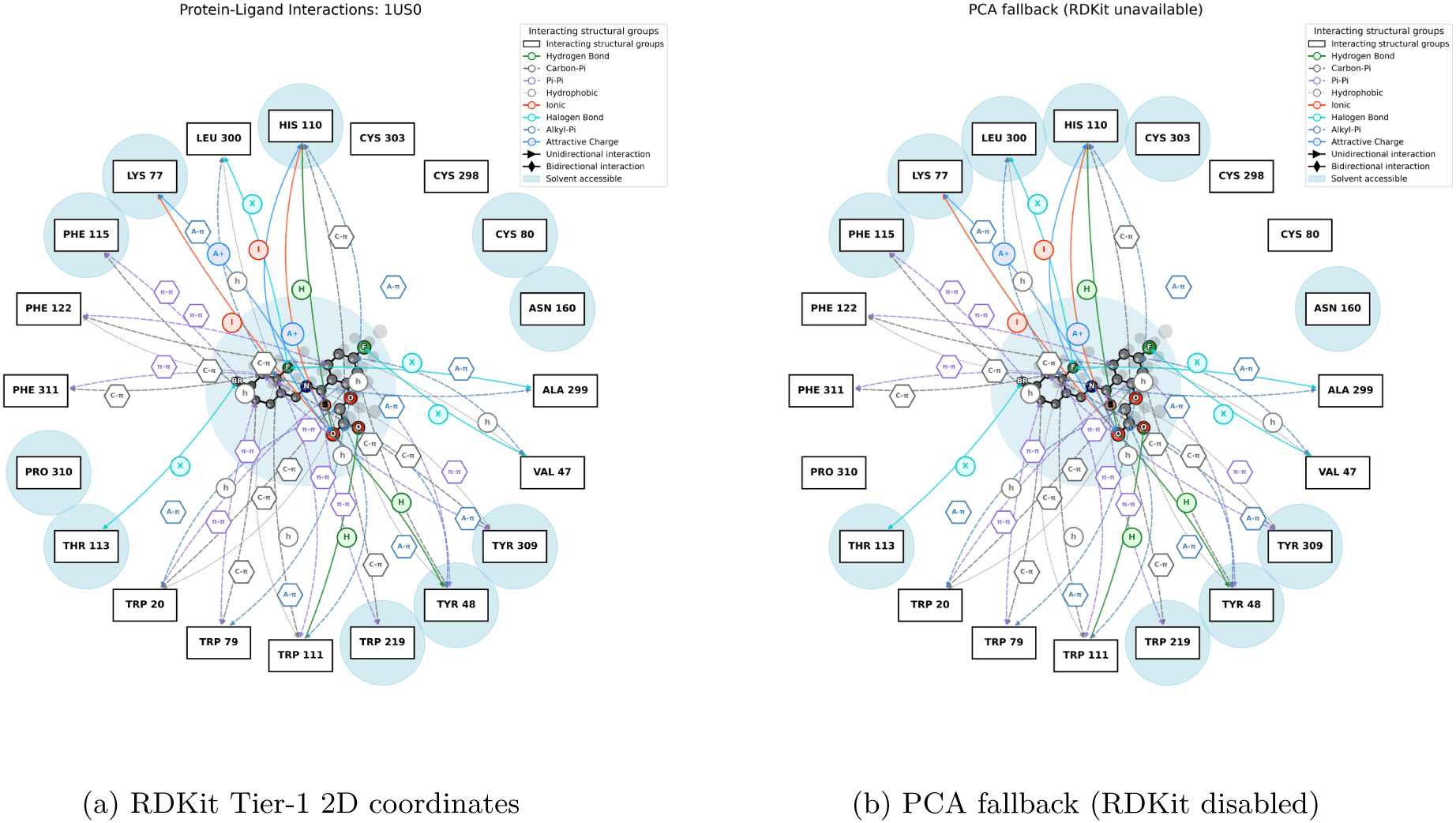
Effect of the two-tier 2D coordinate generation on the aldose reductase–IDD594 diagram (PDB 1US0). (a) With RDKit installed`, Compute2DCoords` lays out the ligand. (b) With RDKit import disabled, the dependency-free PCA projection (Algorithm 2, Tier 2) is used instead. The detected interactions are identical in both panels, including all four halogen bonds and the catalytic hydrogen bonds; only the ligand depiction differs. The PCA projection flattens a 3D conformation onto its two principal axes, so rings are not drawn with regular geometry and atoms can overlap where the molecule folds back on itself. It is a usable fallback rather than an equivalent one, and RDKit is worth installing if the diagram is destined for a figure.

### S1. Installation and System Requirements

PandaMap is distributed via PyPI and installed with a single command: `pip install pandamap`.

**Table S1:**
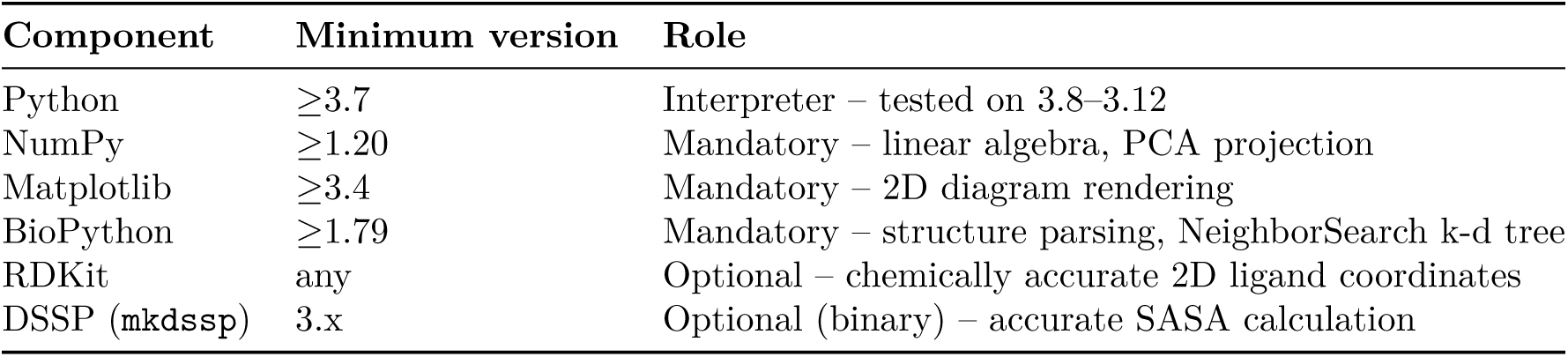
System requirements for PandaMap.

### S2. Command-Line Interface Reference

PandaMap exposes a single entry point, `pandamap:`

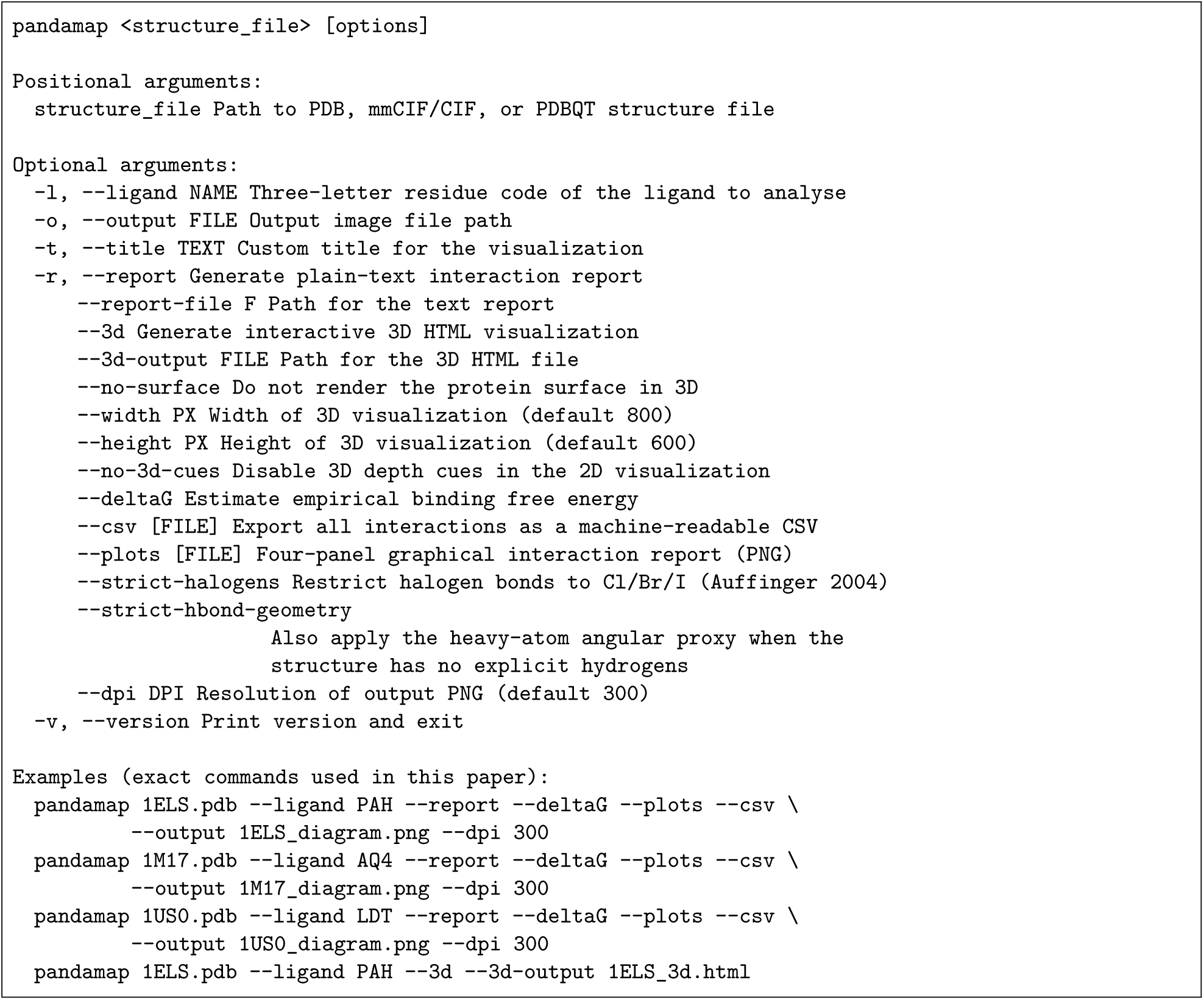

### S3. Python API Reference

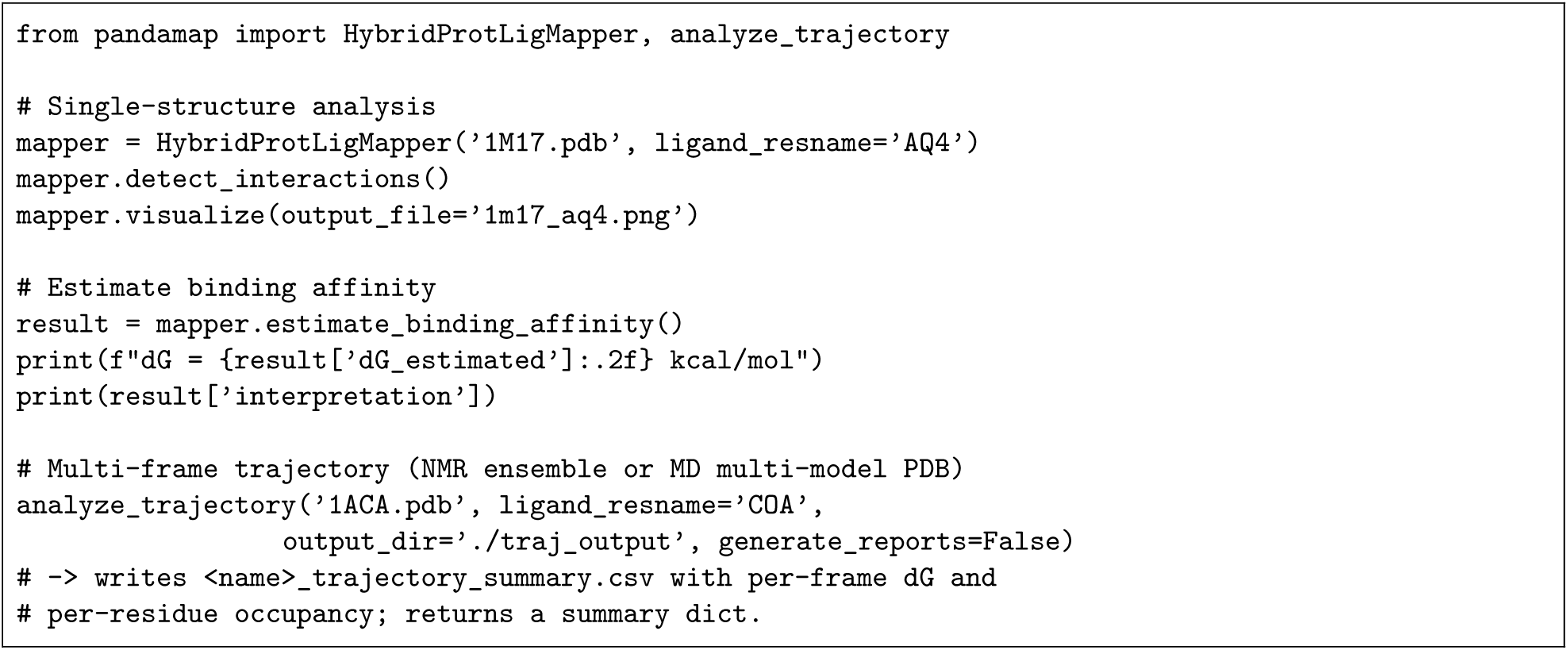

### S4. Protein Aromatic Ring-Atom Name Sets

Only atoms whose names appear below are considered part of an aromatic *π* system; atoms not listed (e.g. C*β*, C*α*) are excluded regardless of residue type, preventing false *π*-system contacts.

**Table S2:**
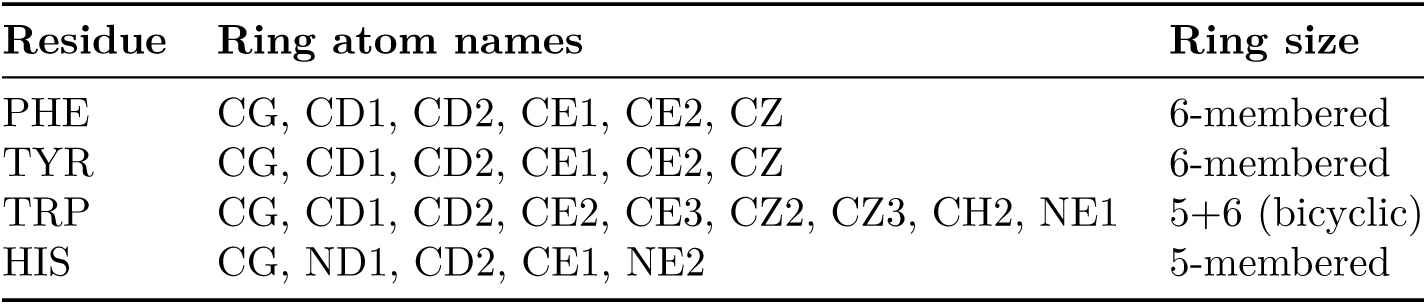
AROMATIC_RING_ATOMS definitions.

### S5. Distance-Cutoff Scientific Justification

**Table S3:**
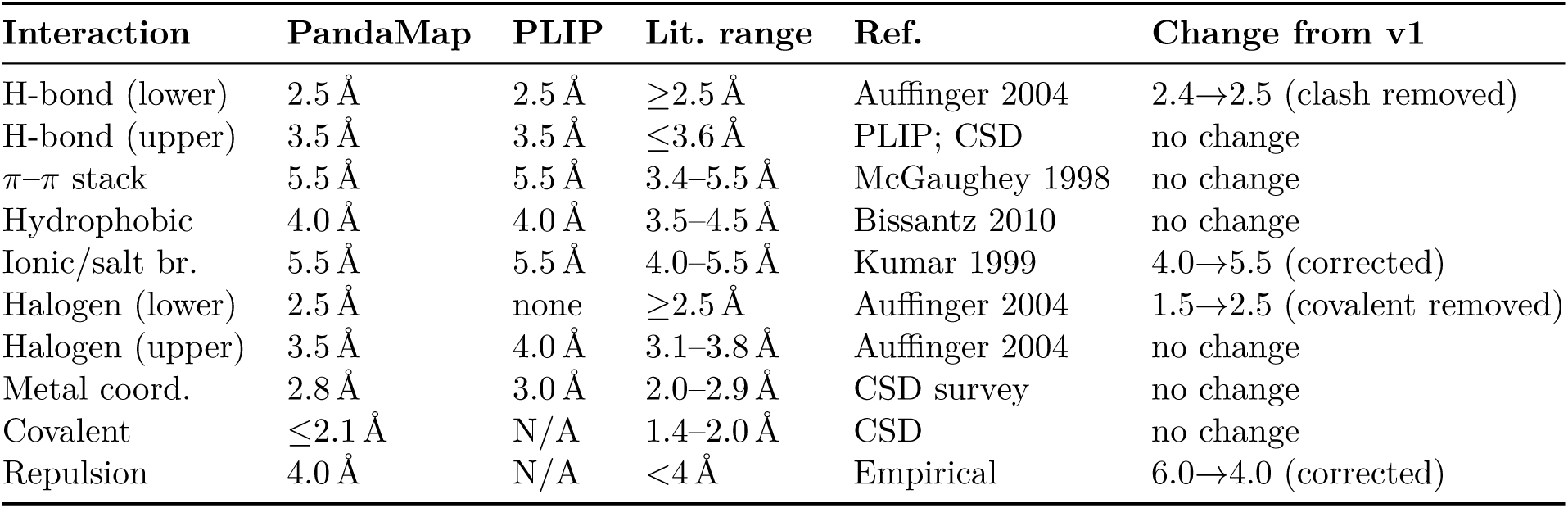
PandaMap distance cutoffs versus PLIP and the literature. Corrections from the original (v1) implementation are marked in the final column. PLIP values are the default thresholds of PLIP 2.3.0; “none” indicates that PLIP applies no lower bound for that class and “N/A” that PLIP does not define the interaction class at all. PLIP additionally imposes angular criteria on halogen bonds that PandaMap does not.

### S6. Empirical Δ*G* Scoring Weights and Parameters

**Table S4:**
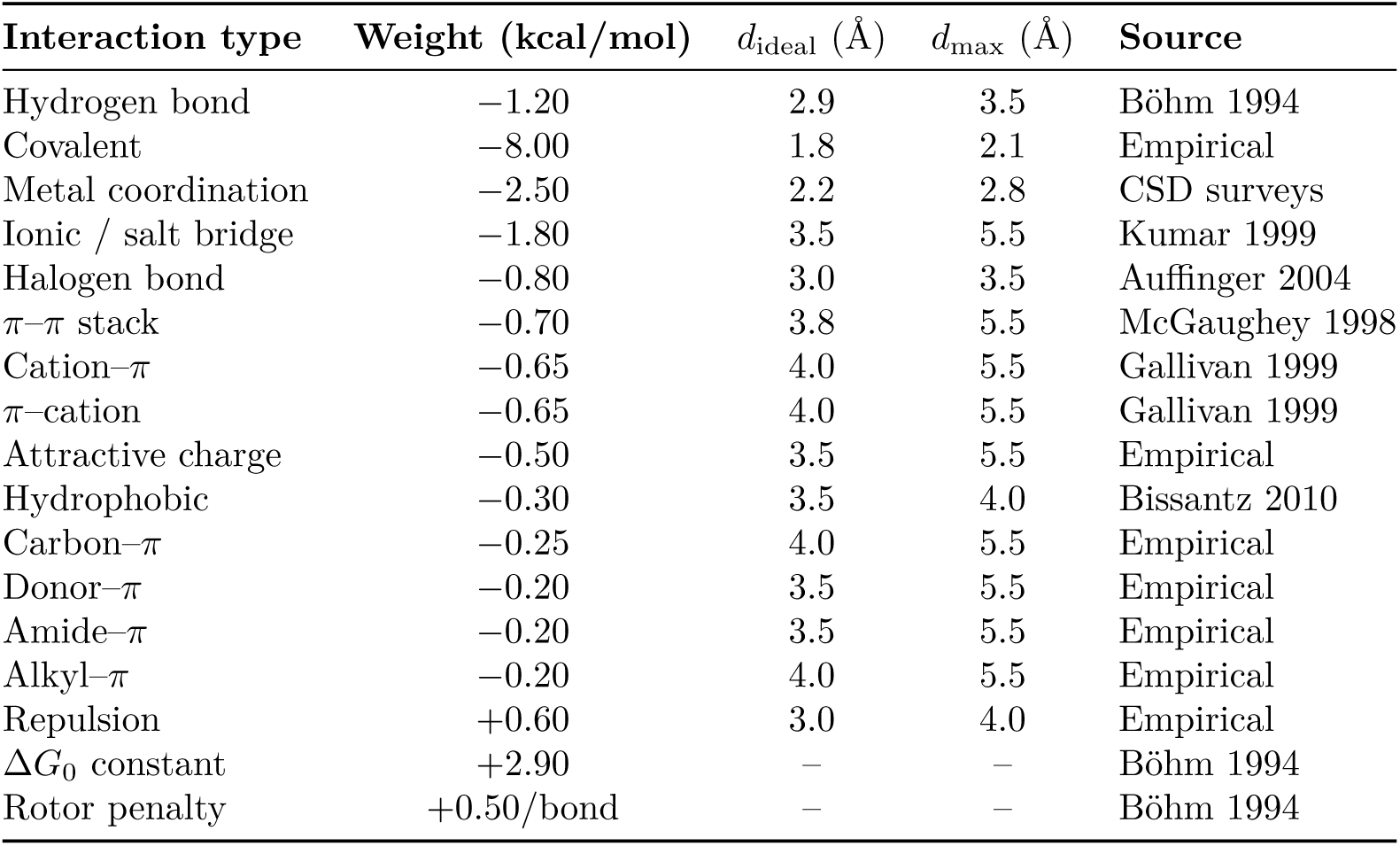
Empirical weights are not regression-fitted to a PDB benchmark; systematic errors of *±*2– 3 kcal/mol are expected. *d*_ideal_ = full-weight distance; *d*_max_ = zero-weight cutoff. In release 4.2.1 the `ionic/salt_bridge` pair and the `cation_pi/pi_cation` pair were not disjoint, so those contacts were weighted twice; both are counted once in 4.3.0 (Section 5). `salt_bridge` shares a detection rule with ionic and no longer carries an independent weight.

| Interaction type | Weight (kcal/mol) | $d_{\text{ideal}}$ (Å) | $d_{\text{max}}$ (Å) | Source |
| --- | --- | --- | --- | --- |
| Hydrogen bond | −1.20 | 2.9 | 3.5 | Böhm 1994 |
| Covalent | −8.00 | 1.8 | 2.1 | Empirical |
| Metal coordination | −2.50 | 2.2 | 2.8 | CSD surveys |
| Ionic / salt bridge | −1.80 | 3.5 | 5.5 | Kumar 1999 |
| Halogen bond | −0.80 | 3.0 | 3.5 | Auffinger 2004 |
| π–π stack | −0.70 | 3.8 | 5.5 | McGaughey 1998 |
| Cation–π | −0.65 | 4.0 | 5.5 | Gallivan 1999 |
| π–cation | −0.65 | 4.0 | 5.5 | Gallivan 1999 |
| Attractive charge | −0.50 | 3.5 | 5.5 | Empirical |
| Hydrophobic | −0.30 | 3.5 | 4.0 | Bissantz 2010 |
| Carbon–π | −0.25 | 4.0 | 5.5 | Empirical |
| Donor–π | −0.20 | 3.5 | 5.5 | Empirical |
| Amide–π | −0.20 | 3.5 | 5.5 | Empirical |
| Alkyl–π | −0.20 | 4.0 | 5.5 | Empirical |
| Repulsion | +0.60 | 3.0 | 4.0 | Empirical |
| $\Delta G_0$ constant | +2.90 | – | – | Böhm 1994 |
| Rotor penalty | +0.50/bond | – | – | Böhm 1994 |

### S7. Behavioural Changes in Release 4.3.0

PandaMap 4.2.x is present on PyPI and may still be in use. The behaviour of several interaction classes changed in 4.3.0, so results are not comparable across the two (Table S5). Most consequentially, halogen-bond detection in 4.2.x matched only fluorine, because the donor test compared element symbols case-sensitively against BioPython’s upper-cased output; chlorine, bromine and iodine were never detected. Charge and *π*-system contacts were also recorded under two keys apiece and so contributed twice to the empirical Δ*G*.

For the benchmark set this moves the enolase estimate from *−*21.97 to *−*18.10 kcal/mol and the EGFR estimate from *−*1.43 to *−*0.86 kcal/mol; the aldose reductase estimate moves the other way, from *−*10.52 to *−*13.92 kcal/mol, because that complex gains a halogen bond and two carboxylate contacts that 4.2.x could not detect. Release 4.3.0 adds a regression suite of 18 tests covering these behaviours.

**Table S5:**
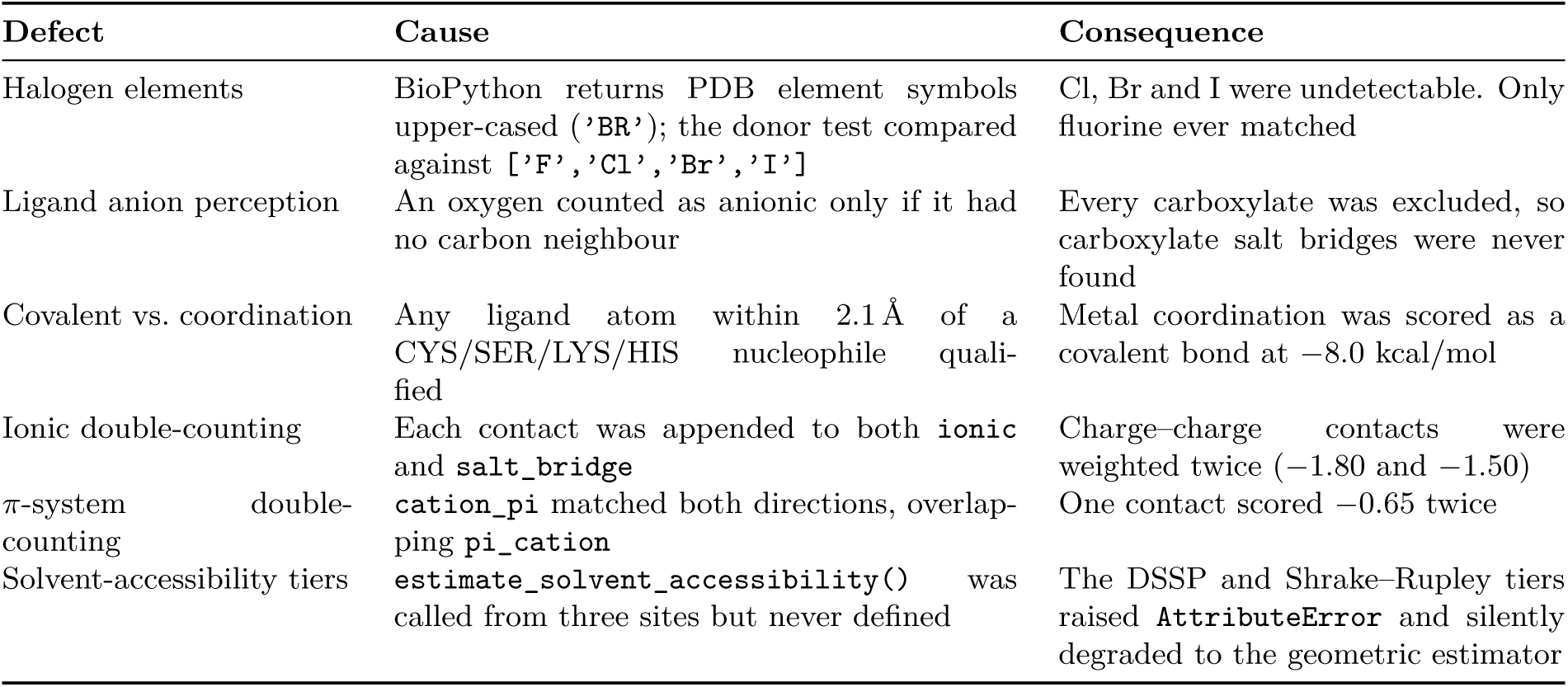
Behavioural changes in release 4.3.0 relative to 4.2.1. Users who ran earlier releases on ligands containing charged groups or halogens should re-run rather than compare results across versions.

### S8. Sample Interaction Report: EGFR–Erlotinib (PDB 1M17)

The following is the verbatim text report produced by `pandamap 1M17.pdb –ligand AQ4 –report`.

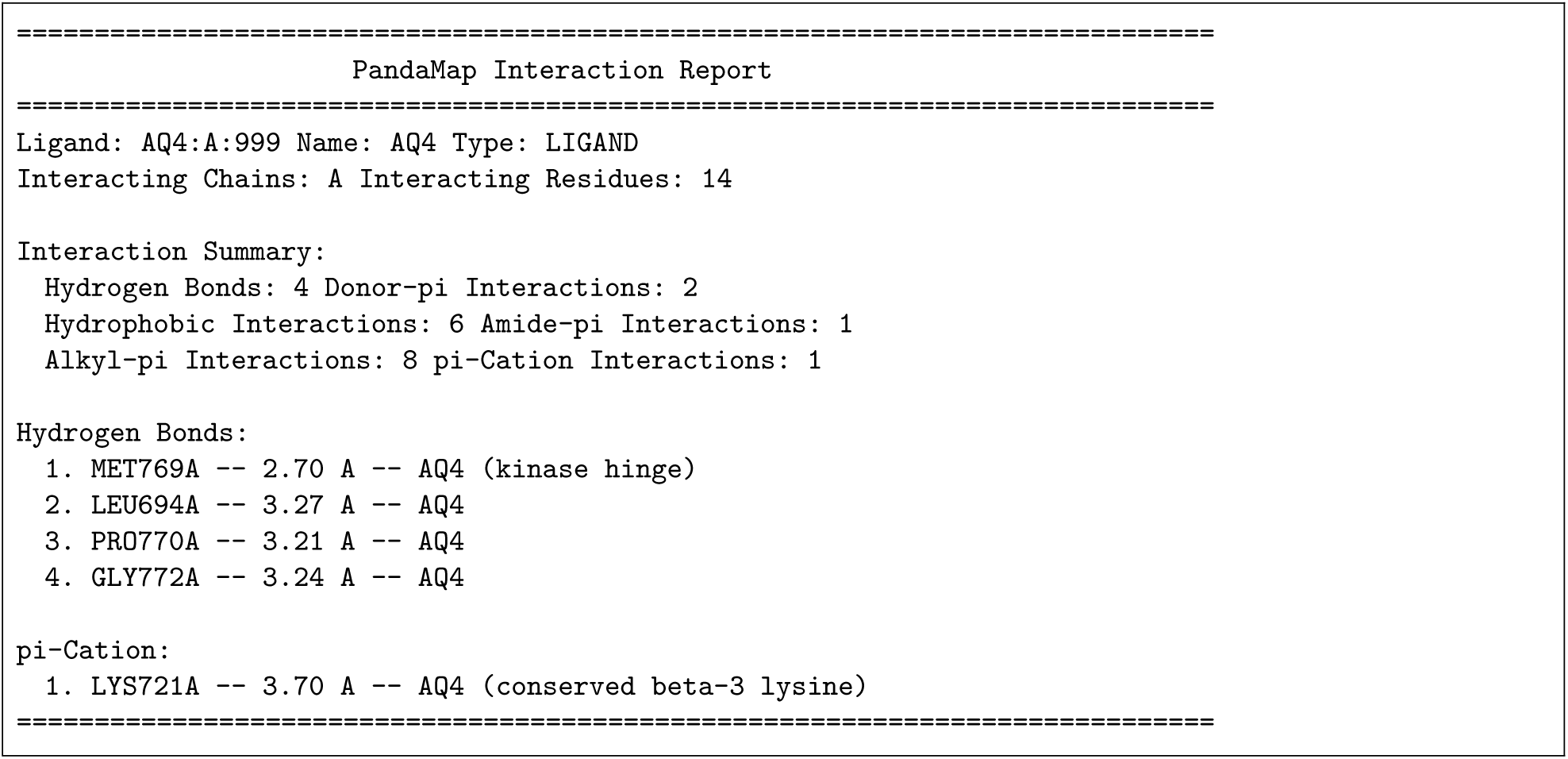

