## Supplementary material for "PandaMap: A Python Package for Comprehensive Visualization of Protein–Ligand Interaction Networks": Figure S1



August 2026 • Software version 4.3.0


---

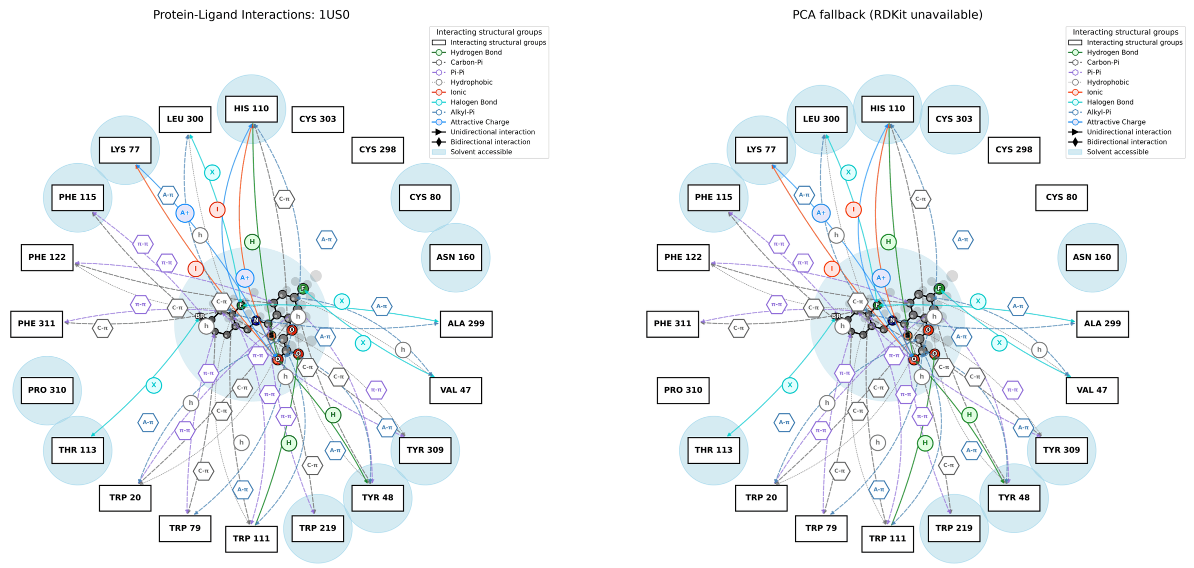


(a) RDKit Tier-1 2D coordinates

(b) PCA fallback (RDKit disabled)

**Figure S1:** Effect of the two-tier 2D coordinate generation on the aldose reductase–IDD594 diagram (PDB 1US0). (a) With RDKit installed, `Compute2DCoords` lays out the ligand. (b) With RDKit import disabled, the dependency-free PCA projection (Algorithm 2, Tier 2) is used instead. The detected interactions are identical in both panels, including all four halogen bonds and the catalytic hydrogen bonds; only the ligand depiction differs. The PCA projection flattens a 3D conformation onto its two principal axes, so rings are not drawn with regular geometry and atoms can overlap where the molecule folds back on itself. It is a usable fallback rather than an equivalent one, and RDKit is worth installing if the diagram is destined for a figure.

### S1. Installation and System Requirements

PandaMap is distributed via PyPI and installed with a single command: `pip install pandamap`.

**Table S1:** System requirements for PandaMap.

| Component | Minimum version | Role |
|---|---|---|
| Python | ≥3.7 | Interpreter – tested on 3.8–3.12 |
| NumPy | ≥1.20 | Mandatory – linear algebra, PCA projection |
| Matplotlib | ≥3.4 | Mandatory – 2D diagram rendering |
| BioPython | ≥1.79 | Mandatory – structure parsing, NeighborSearch k-d tree |
| RDKit | any | Optional – chemically accurate 2D ligand coordinates |
| DSSP (`mkdssp`) | 3.x | Optional (binary) – accurate SASA calculation |

### S2. Command-Line Interface Reference

PandaMap exposes a single entry point, `pandamap`:

```
pandamap <structure_file> [options]

Positional arguments:
  structure_file Path to PDB, mmCIF/CIF, or PDBQT structure file

Optional arguments:
  -l, --ligand NAME Three-letter residue code of the ligand to analyse
  -o, --output FILE Output image file path
  -t, --title TEXT Custom title for the visualization
  -r, --report Generate plain-text interaction report
      --report-file F Path for the text report
      --3d Generate interactive 3D HTML visualization
      --3d-output FILE Path for the 3D HTML file
      --no-surface Do not render the protein surface in 3D
      --width PX Width of 3D visualization (default 800)
      --height PX Height of 3D visualization (default 600)
      --no-3d-cues Disable 3D depth cues in the 2D visualization
      --deltaG Estimate empirical binding free energy
      --csv [FILE] Export all interactions as a machine-readable CSV
      --plots [FILE] Four-panel graphical interaction report (PNG)
      --strict-halogens Restrict halogen bonds to Cl/Br/I (Auffinger 2004)
      --strict-hbond-geometry
                      Also apply the heavy-atom angular proxy when the
                      structure has no explicit hydrogens
      --dpi DPI Resolution of output PNG (default 300)
  -v, --version Print version and exit

Examples (exact commands used in this paper):
  pandamap 1ELS.pdb --ligand PAH --report --deltaG --plots --csv \
          --output 1ELS_diagram.png --dpi 300
  pandamap 1M17.pdb --ligand AQ4 --report --deltaG --plots --csv \
          --output 1M17_diagram.png --dpi 300
  pandamap 1US0.pdb --ligand LDT --report --deltaG --plots --csv \
          --output 1US0_diagram.png --dpi 300
  pandamap 1ELS.pdb --ligand PAH --3d --3d-output 1ELS_3d.html
```

### S3. Python API Reference

```
from pandamap import HybridProtLigMapper, analyze_trajectory
```

```
# Single-structure analysis
mapper = HybridProtLigMapper('1M17.pdb', ligand_resname='AQ4')
mapper.detect_interactions()
mapper.visualize(output_file='1m17_aq4.png')

# Estimate binding affinity
result = mapper.estimate_binding_affinity()
print(f"dG = {result['dG_estimated']:.2f} kcal/mol")
print(result['interpretation'])

# Multi-frame trajectory (NMR ensemble or MD multi-model PDB)
analyze_trajectory('1ACA.pdb', ligand_resname='COA',
                   output_dir='./traj_output', generate_reports=False)
# -> writes <name>_trajectory_summary.csv with per-frame dG and
# per-residue occupancy; returns a summary dict.
```

### S4. Protein Aromatic Ring-Atom Name Sets

Only atoms whose names appear below are considered part of an aromatic $\pi$ system; atoms not listed (e.g. C$\beta$, C$\alpha$) are excluded regardless of residue type, preventing false $\pi$-system contacts.

**Table S2:** `AROMATIC_RING_ATOMS` definitions.

| **Residue** | **Ring atom names** | **Ring size** |
|---|---|---|
| PHE | CG, CD1, CD2, CE1, CE2, CZ | 6-membered |
| TYR | CG, CD1, CD2, CE1, CE2, CZ | 6-membered |
| TRP | CG, CD1, CD2, CE2, CE3, CZ2, CZ3, CH2, NE1 | 5+6 (bicyclic) |
| HIS | CG, ND1, CD2, CE1, NE2 | 5-membered |

### S5. Distance-Cutoff Scientific Justification

**Table S3:** PandaMap distance cutoffs versus PLIP and the literature. Corrections from the original (v1) implementation are marked in the final column. PLIP values are the default thresholds of PLIP 2.3.0; “none” indicates that PLIP applies no lower bound for that class and “N/A” that PLIP does not define the interaction class at all. PLIP additionally imposes angular criteria on halogen bonds that PandaMap does not.

| **Interaction** | **PandaMap** | **PLIP** | **Lit. range** | **Ref.** | **Change from v1** |
|---|---|---|---|---|---|
| H-bond (lower) | 2.5 Å | 2.5 Å | ≥2.5 Å | Auffinger 2004 | 2.4→2.5 (clash removed) |
| H-bond (upper) | 3.5 Å | 3.5 Å | ≤3.6 Å | PLIP; CSD | no change |
| $\pi$–$\pi$ stack | 5.5 Å | 5.5 Å | 3.4–5.5 Å | McGaughey 1998 | no change |
| Hydrophobic | 4.0 Å | 4.0 Å | 3.5–4.5 Å | Bissantz 2010 | no change |
| Ionic/salt br. | 5.5 Å | 5.5 Å | 4.0–5.5 Å | Kumar 1999 | 4.0→5.5 (corrected) |
| Halogen (lower) | 2.5 Å | none | ≥2.5 Å | Auffinger 2004 | 1.5→2.5 (covalent removed) |
| Halogen (upper) | 3.5 Å | 4.0 Å | 3.1–3.8 Å | Auffinger 2004 | no change |
| Metal coord. | 2.8 Å | 3.0 Å | 2.0–2.9 Å | CSD survey | no change |
| Covalent | ≤2.1 Å | N/A | 1.4–2.0 Å | CSD | no change |
| Repulsion | 4.0 Å | N/A | <4 Å | Empirical | 6.0→4.0 (corrected) |

### S6. Empirical $\Delta G$ Scoring Weights and Parameters

**Table S4:** Empirical weights are not regression-fitted to a PDB benchmark; systematic errors of ±2–3 kcal/mol are expected. $d_{\text{ideal}}$ = full-weight distance; $d_{\text{max}}$ = zero-weight cutoff. In release 4.2.1 the `ionic/salt_bridge` pair and the `cation_pi/pi_cation` pair were not disjoint, so those contacts were weighted twice; both are counted once in 4.3.0 (Section S7). `salt_bridge` shares a detection rule with `ionic` and no longer carries an independent weight.

| **Interaction type** | **Weight (kcal/mol)** | $d_{\text{ideal}}$ (Å) | $d_{\text{max}}$ (Å) | **Source** |
|---|---|---|---|---|
| Hydrogen bond | −1.20 | 2.9 | 3.5 | Böhm 1994 |
| Covalent | −8.00 | 1.8 | 2.1 | Empirical |
| Metal coordination | −2.50 | 2.2 | 2.8 | CSD surveys |
| Ionic / salt bridge | −1.80 | 3.5 | 5.5 | Kumar 1999 |
| Halogen bond | −0.80 | 3.0 | 3.5 | Auffinger 2004 |
| $\pi$–$\pi$ stack | −0.70 | 3.8 | 5.5 | McGaughey 1998 |
| Cation–$\pi$ | −0.65 | 4.0 | 5.5 | Gallivan 1999 |
| $\pi$–cation | −0.65 | 4.0 | 5.5 | Gallivan 1999 |
| Attractive charge | −0.50 | 3.5 | 5.5 | Empirical |
| Hydrophobic | −0.30 | 3.5 | 4.0 | Bissantz 2010 |
| Carbon–$\pi$ | −0.25 | 4.0 | 5.5 | Empirical |
| Donor–$\pi$ | −0.20 | 3.5 | 5.5 | Empirical |
| Amide–$\pi$ | −0.20 | 3.5 | 5.5 | Empirical |
| Alkyl–$\pi$ | −0.20 | 4.0 | 5.5 | Empirical |
| Repulsion | +0.60 | 3.0 | 4.0 | Empirical |
| $\Delta G_0$ constant | +2.90 | – | – | Böhm 1994 |
| Rotor penalty | +0.50/bond | – | – | Böhm 1994 |

### S7. Behavioural Changes in Release 4.3.0

PandaMap 4.2.x is present on PyPI and may still be in use. The behaviour of several interaction classes changed in 4.3.0, so results are not comparable across the two (Table S5). Most consequentially, halogen-bond detection in 4.2.x matched only fluorine, because the donor test compared element symbols case-sensitively against BioPython's upper-cased output; chlorine, bromine and iodine were never detected. Charge and $\pi$-system contacts were also recorded under two keys apiece and so contributed twice to the empirical $\Delta G$.

For the benchmark set this moves the enolase estimate from −21.97 to −18.10 kcal/mol and the EGFR estimate from −1.43 to −0.86 kcal/mol; the aldose reductase estimate moves the other way, from −10.52 to −13.92 kcal/mol, because that complex gains a halogen bond and two carboxylate contacts that 4.2.x could not detect. Release 4.3.0 adds a regression suite of 18 tests covering these behaviours.

**Table S5:** Behavioural changes in release 4.3.0 relative to 4.2.1. Users who ran earlier releases on ligands containing charged groups or halogens should re-run rather than compare results across versions.

| Defect | Cause | Consequence |
|---|---|---|
| Halogen elements | BioPython returns PDB element symbols upper-cased (`'BR'`); the donor test compared against `['F','Cl','Br','I']` | Cl, Br and I were undetectable. Only fluorine ever matched |
| Ligand anion perception | An oxygen counted as anionic only if it had no carbon neighbour | Every carboxylate was excluded, so carboxylate salt bridges were never found |
| Covalent vs. coordination | Any ligand atom within 2.1 Å of a CYS/SER/LYS/HIS nucleophile qualified | Metal coordination was scored as a covalent bond at −8.0 kcal/mol |
| Ionic double-counting | Each contact was appended to both `ionic` and `salt_bridge` | Charge–charge contacts were weighted twice (−1.80 and −1.50) |
| $\pi$-system double-counting | `cation_pi` matched both directions, overlapping `pi_cation` | One contact scored −0.65 twice |
| Solvent-accessibility tiers | `estimate_solvent_accessibility()` was called from three sites but never defined | The DSSP and Shrake–Rupley tiers raised `AttributeError` and silently degraded to the geometric estimator |

### S8. Sample Interaction Report: EGFR–Erlotinib (PDB 1M17)

The following is the verbatim text report produced by `pandamap 1M17.pdb -ligand AQ4 -report`.

```
================================================================================
                     PandaMap Interaction Report
================================================================================
Ligand: AQ4:A:999 Name: AQ4 Type: LIGAND
Interacting Chains: A Interacting Residues: 14

Interaction Summary:
  Hydrogen Bonds: 4 Donor-pi Interactions: 2
  Hydrophobic Interactions: 6 Amide-pi Interactions: 1
  Alkyl-pi Interactions: 8 pi-Cation Interactions: 1

Hydrogen Bonds:
  1. MET769A -- 2.70 A -- AQ4 (kinase hinge)
  2. LEU694A -- 3.27 A -- AQ4
  3. PRO770A -- 3.21 A -- AQ4
  4. GLY772A -- 3.24 A -- AQ4

pi-Cation:
  1. LYS721A -- 3.70 A -- AQ4 (conserved beta-3 lysine)
================================================================================
```
